# Mesolimbic microglia display regionally distinct developmental trajectories

**DOI:** 10.1101/764233

**Authors:** KT Hope, IA Hawes, A Bonci, LM De Biase

**Affiliations:** Department of Physiology, David Geffen School of Medicine, University of California Los Angeles, Los Angeles, CA 90095, USA; Intramural Research Program, National Institute on Drug Abuse, National Institutes of Health, Baltimore MD, 21224, USA; Solomon H. Snyder Department of Neuroscience, Johns Hopkins University School of Medicine, Baltimore MD 21205, USA; Department of Psychiatry and Behavioral Sciences, Johns Hopkins University School of Medicine, Baltimore, MD 21287, USA

**Author notes:** These authors contributed equally to this work (KT Hope, IA Hawes). Current contact; Global Institutes on Addiction, Miami, FL, 33131, USA. Co-senior authorship (A Bonci, LM De Biase). **Correspondence:** Dr. Lindsay De Biase, 10833 Le Conte Ave, CHS 77-200A, Los Angeles, CA 90095.

**Keywords:** mouse, ventral tegmental area, nucleus accumbens, cell density, programmed cell death, lysosome, phagocytosis

## Abstract

Microglia play critical roles during CNS development and undergo dramatic changes in tissue distribution, morphology, and gene expression as they transition from embryonic to neonatal to adult microglial phenotypes. Despite the magnitude of these phenotypic shifts, little is known about the time course and dynamics of these transitions and whether they vary across brain regions. Here we define the time course of microglial maturation in key regions of the basal ganglia in mice where significant regional differences in microglial phenotype are present in adults. We found that microglial density peaks in the ventral tegmental area (VTA) and nucleus accumbens (NAc) during the third postnatal week, driven by a burst of microglial proliferation. Microglial abundance is then refined to adult levels through a combination of tissue expansion and microglial programmed cell death. This overproduction and refinement of microglia was significantly more pronounced in the NAc and was accompanied by a sharp peak in NAc microglial lysosome abundance in the third postnatal week. Collectively, these data identify a key developmental window when elevated microglial density in discrete basal ganglia nuclei may support circuit refinement and could increase susceptibility to inflammatory insults.

## INTRODUCTION

In addition to their well-recognized roles in CNS immune surveillance and injury responses, microglia have now been shown to execute critical functions during embryonic and neonatal development. During embryogenesis, microglia promote maturation of the vasculature, support survival and regulate programmed cell death of specific neuronal populations, and shape axon guidance during circuit wiring (Reemst *et al.*, 2016; Mosser *et al.*, 2017). During neonatal periods, microglia continue to shape composition of cell populations and participate in synapse formation, synapse elimination, and maturation of synaptic function (Frost & Schafer, 2016; VanRyzin *et al.*, 2018), with critical consequences for sensory function and brain sexual dimorphism. Consistent with the idea that these cells play critical roles in CNS development, microglia have been implicated in the pathogenesis of multiple neurodevelopmental disorders (Tay *et al.*, 2018).

In contrast to other major glial cell populations, microglia arise from primitive macrophage progenitors in the yolk sac (Ginhoux *et al.*, 2013). These progenitors invade the developing neuroectoderm during early embryogeneis and then go on to migrate, proliferate, and colonize the CNS with the mature resident microglial cell population. During embryonic and neonatal periods microglia are unevenly distributed throughout the CNS, clustering near the ventricles, pial surface, and along axonal tracts (Arnoux *et al.*, 2013; Squarzoni *et al.*, 2014; Reemst *et al.*, 2016; Mosser *et al.*, 2017). In contrast, in the mature CNS, these cells are tiled throughout the brain. Transcriptome studies indicate that these shifts in microglial tissue distribution and morphology are accompanied by massive changes in gene expression that suggest discrete steps in microglial maturation between embryonic, neonatal, and adult phenotypes (Butovsky *et al.*, 2014; Bennett *et al.*, 2016; Matcovitch-Natan *et al.*, 2016; Hammond *et al.*, 2019; Li *et al.*, 2019). However, the dynamics of these transitions in tissue distribution, morphology, and phenotype have not been fully mapped out, particularly the transition from neonatal to adult phenotypes. Whether the time course and mechanisms underlying these phenotypic transitions are equivalent throughout the CNS also remains unknown (Mosser *et al.*, 2017).

Several neurodevelopmental disorders in which microglia been implicated involve dysfunction of mesolimbic dopamine signaling, such as schizophrenia and ADHD (Tay *et al.*, 2018), suggesting that detailed information about microglial development in these brain regions could provide new insight into windows of susceptibility and pathological mechanisms. However, developmental trajectories of microglia in these brain regions have not been defined. Our previous studies found that microglia within the mature basal ganglia exhibit region-specific phenotypes and that these regional differences emerge during early postnatal development (De Biase *et al.*, 2017). This pronounced regional variation in adult microglia raises questions about whether the dynamics of microglial maturation and regulatory factors involved may vary significantly.

Here we define the time course over which microglia colonize the developing mesolimbic system and acquire their mature distribution and morphology. These studies provide a critical foundation for better understanding how these cells contribute to circuit maturation within the mesolimbic system and identify regions and temporal windows during which perturbation of microglial cells is likely to contribute to neurodevelopmental disorders that involve these circuits.

## MATERIALS AND METHODS

### Transgenic Mice

Transgenic mice expressing enhanced green fluorescent protein (EGFP) under control of the microglial specific promoter (CX3CR1)(Jung *et al.*, 2000) were used to visualize the detailed morphology and density of microglial cells. *CX3CR1*^*EGFP / EGFP*^ breeders were originally obtained from Jackson labs (Stock # 005582). In these mice EGFP is knocked into the CX3CR1 locus. All mice used for experiments were heterozygous for the transgene (*CX3CR1*^*EGFP/+*^). C57Bl6 wildtype mice were used for comparison in some experiments.

In all experiments, both male and female mice were used, and the number of males and females in each analysis group was balanced. Mice were housed in normal light dark cycle and had ad libidum access to food and water. All experiments were performed in strict accordance with protocols approved by the Animal Care and Use Committees at NIDA and UCLA (NIDA protocol #18-OSD-6 and UCLA protocol #18-103).

### Tissue collection and immunohistochemistry

Tissue from *CX3CR1*^*EGFP / +*^ or C57Bl6 wildtype mice was collected between postnatal days (P) P8 and P60. Mice were anesthetized with pentobarbital (100 mg/kg). Microglia show circadian-based changes in morphology (Hayashi *et al.*, 2013); all perfusions for this study were performed between 9:00am-12:00pm. Mice were perfused transcardially with 1M phosphate buffered saline (PBS) followed by 4% paraformaldehyde (PFA) in 1M PBS. The brain was isolated and placed in 4% PFA for an additional 4 hours for post-fixation at 4^°^C and was then transferred to 1M PBS with 0.1% NaAz.

Brains were sectioned on a vibratome at a thickness of 60 microns (μm) or cryoprotected in 30% sucrose, frozen and embedded in OTC medium, and sectioned on a cryostat at a thickness of 50 μm. Free floating sections were permeabilized/blocked with a solution of 5% Normal Donkey Serum (NDS) and 0.3% Triton X-100 in 1M PBS for 2 hours with rotation at room temperature (RT). Sections were incubated with primary antibodies prepared in 5% Normal Donkey Serum and 0.05% Triton X-100 in 1M PBS with rotation overnight at 4^°^C. Sections were incubated with secondary antibodies prepared in 5% NDS in 1M PBS with rotation for 2 hours at RT. Control sections incubated with secondary antibody alone did not result in labeling of cells. Primary antibodies used include: chicken α GFP (Aves, 1:1000), rabbit α Iba1 (Wako, 1:500), rabbit α caspase 3 (Cell Signaling, 1:200), mouse α Ki67 (Abcam, 1:500), and mouse α Tyrosine Hydroxylase (Sigma, 1:5000). Secondary antibodies used include: Alexa 488, Alexa 594 and Alexa 647 fluorophore conjugated antibodies against Rabbit, Chicken, Mouse, or Guinea Pig. (1:500; all raised in donkey; Jackson ImmunoResearch).

### Image acquisition and analysis

For measurement of microglial cell density images were acquired with an Olympus MXV10 epifluorescence microscope and then imported into ImageJ software (NIH) for analysis. For analysis of developmental tissue expansion, images were acquired on a Zeiss Apotome epifluorescence microscope and brain nuclei were manually outlined in ImageJ using standard anatomical landmarks. For analysis of microglial cell branching, Ki67 expression, capspase 3 expression, and microglial lysosome content images were acquired on an Olympus FV1000 confocal fluorescence microscope. Stacks of confocal images (z-stacks) with a z interval of 1.5 μm were taken and imported into Image J software. For analysis of microglial tissue coverage, mean pixel intensity of the dimmest cell processes was measured at 10-15 locations diagonally across each image. The average of these values was taken as a threshold for determining the % of pixels above (representing microglial cell somas and processes) and below this threshold. Within the NAc, analyzed images were acquired at the boundary between core and shell (identified anatomically). In the VTA, analyzed images were medial to the medial lemniscus and included the parabrachial pigmented area, as well as portions of the parafasciculus retroflexus area and paranigral nucleus. For each time point and brain region, 2-3 images per mouse were acquired for 3-6 different mice.

### Statistical comparisons

All graphed values are shown as mean ± SEM. Statistical details of experiments (statistical tests used, exact value of n, what n represents) can be found in the results and figures legends. In general, statistical significance was assessed using one- or two-way ANOVA (Figures 1C, 1D, 1E, 2B, 3C, 3D, 3E, 4B, 6D, 6E, 6F, 7B, 7C, S1B, S1C, S2, S5A, S5B). Posthoc comparisons were carried out using student’s t-test with Bonferroni correction for multiple comparisons. Data are sufficiently normal and variance within groups is sufficiently similar for use of parametric tests. Paired t-tests were used to compare trends across two different regions within individual mice (Figures 6G, S3B). Linear regression tests were used to test the dependence of variable *y* on variable *x* (Figure S3C).

**Figure 1.**
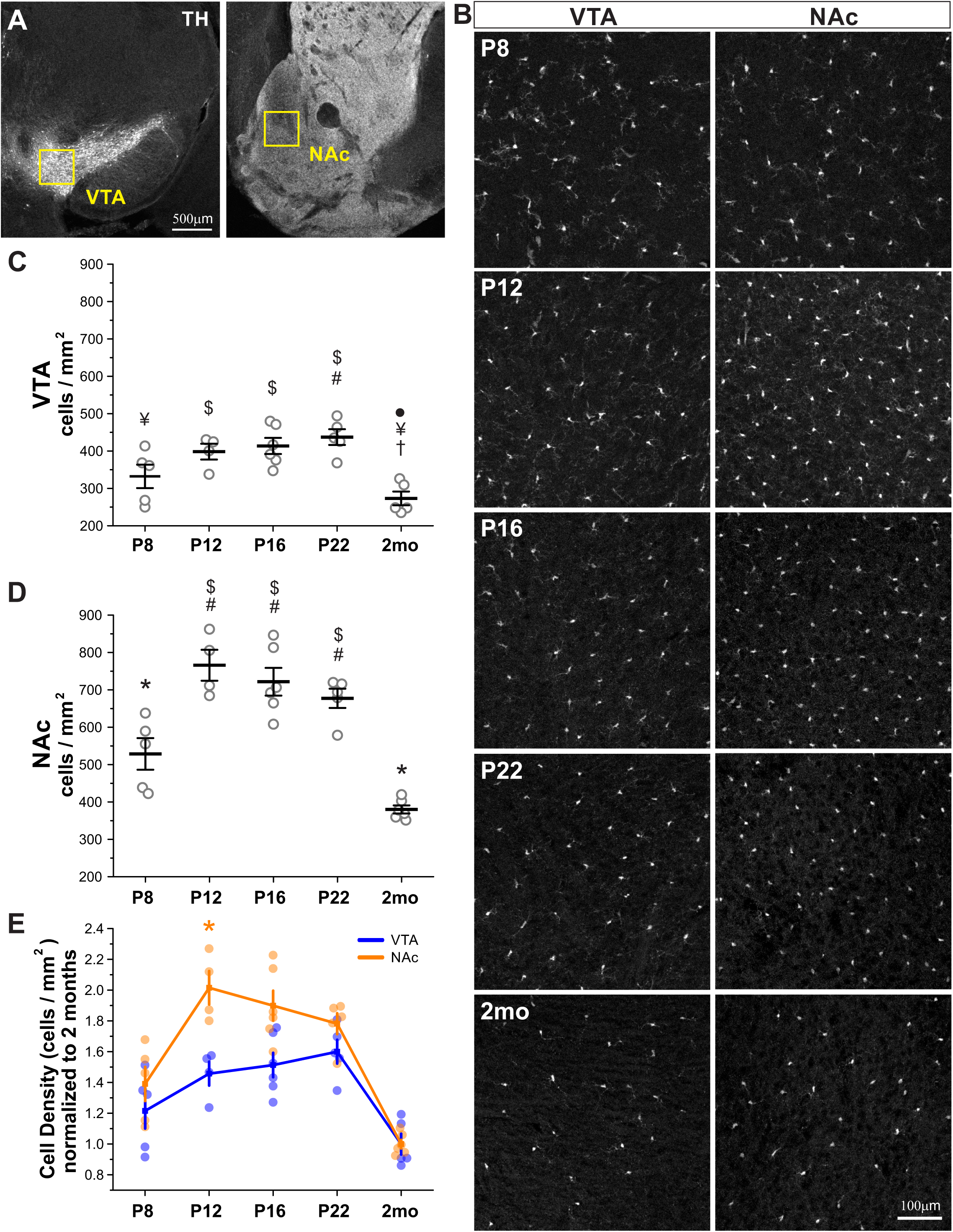
Pronounced overproduction of nucleus accumbens microglia during postnatal development. **(A)** Coronal brain sections from a *CX3CR1*^*EGFP/+*^ mouse immunostained for tyrosine hydroxylase (TH), which labels cell bodies and projections of dopamine neurons. *Yellow boxes* indicate the location of images acquired for analysis in the ventral tegmental area (VTA) and the nucleus accumbens (NAc). **(B)** Representative images showing the distribution of EGFP+ microglia at postnatal days (P) 8,12,16,22, and 2 months (2mo) in the VTA and NAc. **(C)** Quantification of microglial density during early postnatal development in the VTA. Each data point depicts the average density value obtained for one mouse (2-3 images analyzed per mouse). n=4-6 mice analyzed per region. ANOVA F(_4,24_) = 8.3, P = 0.0004. # P < 0.05 vs. P8, • P < 0.05 vs. P12, † P < 0.05 vs. P16, ¥ P < 0.05 vs. P22, $ P < 0.05 vs. 2mo. **(D)** Quantification of microglial density during early postnatal development in the NAc. Each data point depicts the average density value obtained for one mouse (2-3 images analyzed per mouse). n=4-6 mice analyzed per region. ANOVA F(_4,25_) = 25.1, P < 0.00001. # P < 0.05 vs. P8, $ P < 0.05 vs. 2mo. * P < 0.05 all comparisons. **(E)** Increases and decreases in microglial density in the VTA and NAc normalized to the mean density attained in each region at 2mo. 2-way ANOVA; main effect of age, F(_4,41_) = 29.7, P < 0.00001; main effect of brain region, F(_1,41_) = 22.8, P <0.00001; age x brain region interaction, F(_5,41_) = 3.0, P = 0.03. * P = 0.009 P12 VTA vs. NAc posthoc comparison.

## RESULTS

### Density of mesolimbic microglia peaks during early postnatal development

In young adult mice, microglial density varies dramatically across distinct nuclei of the basal ganglia (De Biase *et al.*, 2017). These differences emerge early in postnatal development (De Biase *et al.*, 2017), but the detailed time course over which mesolimbic microglia achieve their mature density and distribution has not been described. To analyze microglial maturation within the developing mesolimbic system, we quantified microglial cell density within two key interconnected basal ganglia nuclei, the ventral tegmental area (VTA) and its projection target, the nucleus accumbens (NAc), at postnatal day (P) 8, 12, 16, 22, and 2 months using transgenic mice that express EGFP under control of the endogenous fractalkine promoter (*CX3CR1*^*EGFP/+*^ mice)(**Fig. 1A,B)**. In the VTA of P8 mice, microglia were evenly distributed and cell density exceeded the 205 ± 24 cells / mm^2^ reported previously for the same brain region at P6 (**Fig. 1B,C**)(De Biase *et al.*, 2017). Between P8-P22, microglial density in the VTA continued to increase at a steady rate reaching a maximum density of 437 ± 21 cells / mm^2^ at P22, roughly double that reported at P6. Between P22 and 2 months, microglial density fell significantly to reach 273 ± 10 cells / mm^2^, consistent with previous reports of 262 ± 10 cells / mm^2^ for this brain region in young adult mice.

In the NAc of P8 mice, microglia were also evenly distributed and, similar to the VTA, cell density already exceeded the 220 ± 19 cells / mm^2^ reported previously for the NAc at P6 (**Fig. 1D**)(De Biase *et al.*, 2017). In contrast to the VTA, however, the increase in microglial density at P8 represented a nearly 3-fold elevation over P6 microglial numbers. Between P8 and P12, NAc microglial density nearly doubled again, reaching a peak of 757 ± 41 cells / mm^2^. Rather than continuing to increase as observed in the VTA, microglial numbers then remained steady or even began to decline between P12 and P22. Between P22 and 2 months, density of NAc microglia fell sharply to 380 ± 11 cells / mm^2^, consistent with previous reports of 380 ± 7 cells / mm^2^ for this brain region in young adult mice.

Collectively, these results indicate that there is a pronounced increase in microglial cell numbers in multiple mesolimbic nuclei during the first 3 postnatal weeks, with peak densities significantly exceeding those observed in adult mice. The apparent differences between VTA and NAc in the magnitude and time course of this microglial density increase and subsequent reduction was significant. Two-way ANOVA analysis of changes in microglial density during postnatal development revealed significant main effects of age (P < 0.00001), brain region (P <0.00001), and a significant interaction between the two (P = 0.0006). Normalizing densities of microglia at early postnatal timepoints to the density found in each brain region in young adult mice (2 mo mice), emphasized the steeper increase in microglial density within the NAc (**Fig. 1E**, 2-way ANOVA; main effect of age, F(_4,41_) = 29.7, P < 0.00001; main effect of brain region, F(_1,41_) = 22.8, P <0.00001; age x brain region interaction, F(_5,41_) = 3.0, P = 0.03).

This time course of increasing and subsequently declining microglial density was also observed in wildtype mice (**Fig. S1 A,B**). In wildtype mice, the magnitude of microglial overproduction was also more pronounced in the NAc as compared to the VTA (**Fig. S1 C**, 2-way ANOVA; main effect of age, F(_4,20_) = 106.6, P < 0.00001; main effect of brain region, F(_1,20_) = 202.5, P <0.00001; age x brain region interaction, F(_5,20_) = 20.2, P <0.00001). Collectively, these findings indicate that microglia do not follow the same program of maturation across different brain regions and raise questions about the underlying regulatory factors driving these differences. These findings also highlight the particularly elevated density of microglia in the NAc during the second and third postnatal weeks.

### Microglial tissue coverage increases linearly over postnatal development

During the first postnatal week, microglia exhibit amoeboid morphologies and limited process ramification (Schafer *et al.*, 2012; Lenz *et al.*, 2013; De Biase *et al.*, 2017). Elaboration of highly ramified processes then proceeds over subsequent postnatal weeks. To examine the morphological maturation of microglial cells in the mesolimbic system, we used *CX3CR1*^*EGFP/+*^ mice to visualize the fine branching structure of these cells in the VTA and NAc at different developmental time points. In both the NAc and VTA, microglial process branching is limited at P8 and some cells with amoeboid morphology can still be observed (**Fig. 2A**). By P12, most cells in both the NAc and VTA exhibit extensive process branching and by P22, cell morphology in both regions closely resembles that in the adult.

**Figure 2.**
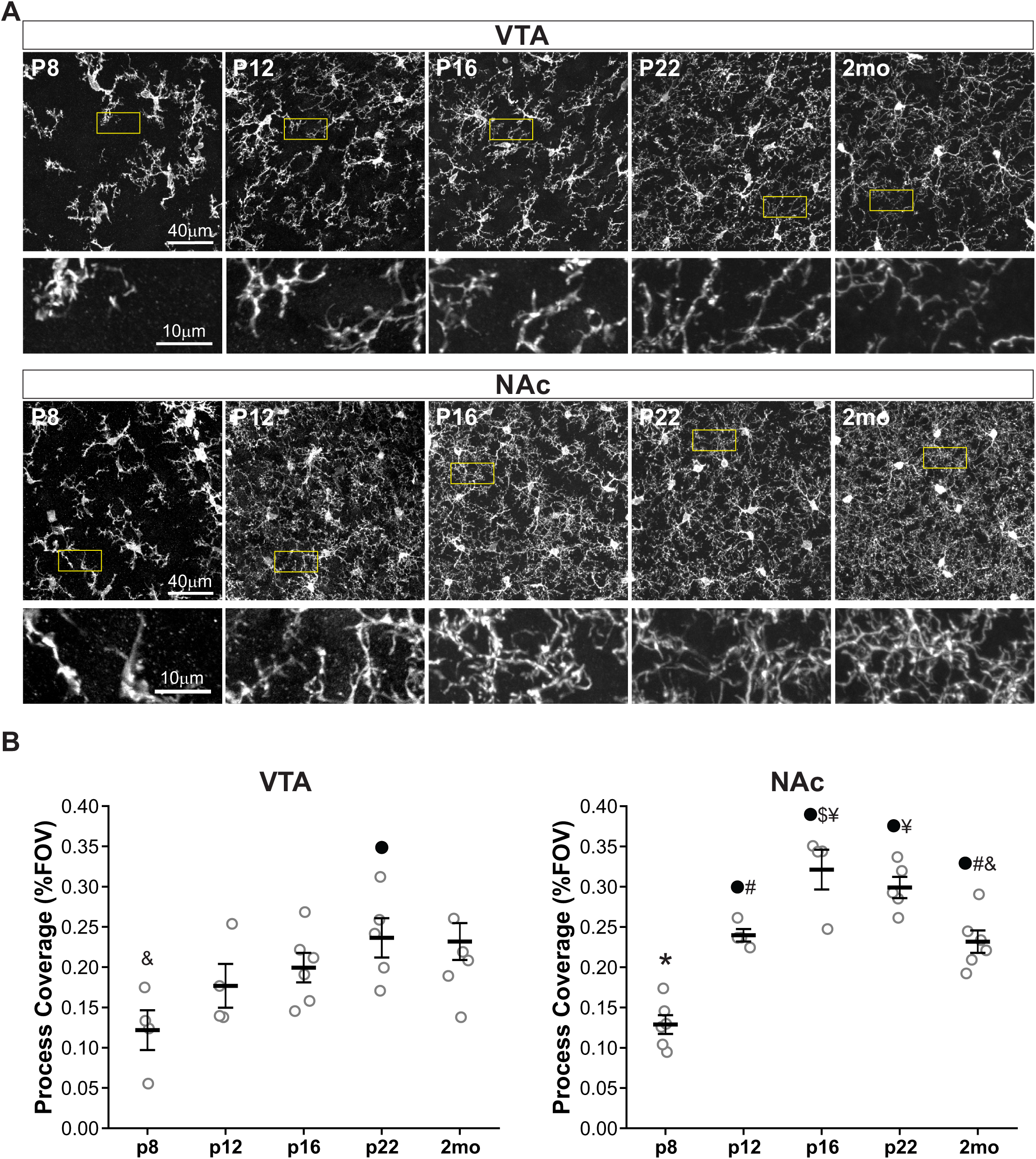
Overproduction of NAc microglial processes during postnatal development. **(A)** Representative high magnification images of microglia in the VTA and NAc during early postnatal development. Regions highlighted by *yellow boxes* enlarged in lower panels to show microglial branching patterns. **(B)** Quantification of VTA microglial tissue coverage, calculated as the percent coverage of the field of view by microglial processes (not including somas). ANOVA F(_4,19_) = 3.2, p=.03521. • P < 0.05 vs. P8, & P < 0.05 vs. P22. **(C)** Quantification of NAc microglial tissue coverage, calculated as the percent coverage of the field of view by microglial processes (not including somas). ANOVA F(_4,20_) = 27.5, P < 0.00001. * P < 0.001 vs. all comparisons, P < 0.001 vs. P8, $ P < 0.05 vs. P12, # P < 0.05 vs. P16, & P < 0.05 vs. P22, ¥ P < 0.05 vs. 2mo. Data points depict average values obtained for one mouse (2-3 images analyzed per mouse); n=3-6 mice analyzed per timepoint per region.

To quantify these developmental changes, we analyzed the degree to which microglial processes cover a particular field of view (% FOV covered by microglial processes). Microglial process coverage in the VTA increased steadily between P8 and P22 (**Fig. 2B**) after which process coverage remained comparable at 2 months. In contrast, in the NAc, microglial process coverage increased sharply between P8 and P12 and again between P12 and P16, which represented a peak in microglial process coverage in the NAc (**Fig. 2B**). By 2 months, microglial process coverage had decreased significantly from this peak level. Similar results were obtained when taking both microglial processes and somas into account for calculations of tissue coverage (**Fig. S2**). These apparent differences between VTA and NAc in timecourse and magnitude of process coverage were significant; two-way ANOVA analysis of changes in microglial process coverage during postnatal development revealed significant main effects of age (F(_1,39_) = 18.0, P <0.00001), brain region (F(_1,39_) = 22.0, P = 0.00003), and a significant interaction between the two (F(_5,39_) = 2.6, P = 0.048). Collectively, these observations reinforce the idea that VTA and NAc microglia follow distinct programs of maturation and again identify the second and third postnatal weeks as a unique window of elevated microglial occupancy within the NAc.

### Proliferation of resident microglia at P8 contributes to increase in cellular density

Lineage tracing studies have indicated that CNS resident microglia all derive from progenitors that invade the CNS during late embryogenesis (Ginhoux *et al.*, 2010, 2013), suggesting that these progenitors will need to proliferate substantially in order to increase the microglial population from neonatal abundance to mature numbers. Indeed, recent whole transcriptome analyses of microglia in development suggest that a substantial proportion of the microglial population is proliferative in neonatal periods but that this has ceased by P30 (Hammond *et al.*, 2019). To investigate whether proliferation of resident microglia underlies increases in VTA and NAc microglial cell density from P8 to P12, we examined Ki67 immunostaining in these brain regions. Both EGFP^+^Ki67^+^ and EGFP^-^Ki67^+^ proliferating cells were readily apparent in the early postnatal VTA and NAc (**Fig. 3A** and **Fig. S3**). Some Ki67^+^ microglia appeared isolated, whereas others showed clear morphological profiles of cell division and distinct nuclei in daughter cells (**Fig. 3B** and **Fig. S3A**). Quantitative analysis revealed that the density of EGFP^+^Ki67^+^ was highest at P8 and fell off rapidly after P12 (**Fig. 3C**), consistent with achievement of peak cell density by this age. Examination of the proportion of proliferating microglia revealed a higher percentage of EGFP^+^Ki67^+^ cells in P8 and P12 VTA compared to the NAc (**Fig. 3D**; 2-way ANOVA; main effect of age, F(_4,42_) = 6.0, P = 0.0007; main effect of brain region, F(_1,42_) = 5.1, P = 0.03; age x brain region interaction, F(_5,42_) = 1.8, P = 0.14.), suggesting that proliferation falls off more slowly in the VTA. This would be consistent with the slightly later peak in microglial density in the VTA compared to the NAc. Consistent with this idea, within individual mice at P8-12, the proportion of proliferating microglia was higher in the VTA than in the NAc (**Fig. S3 B**).

**Figure 3.**
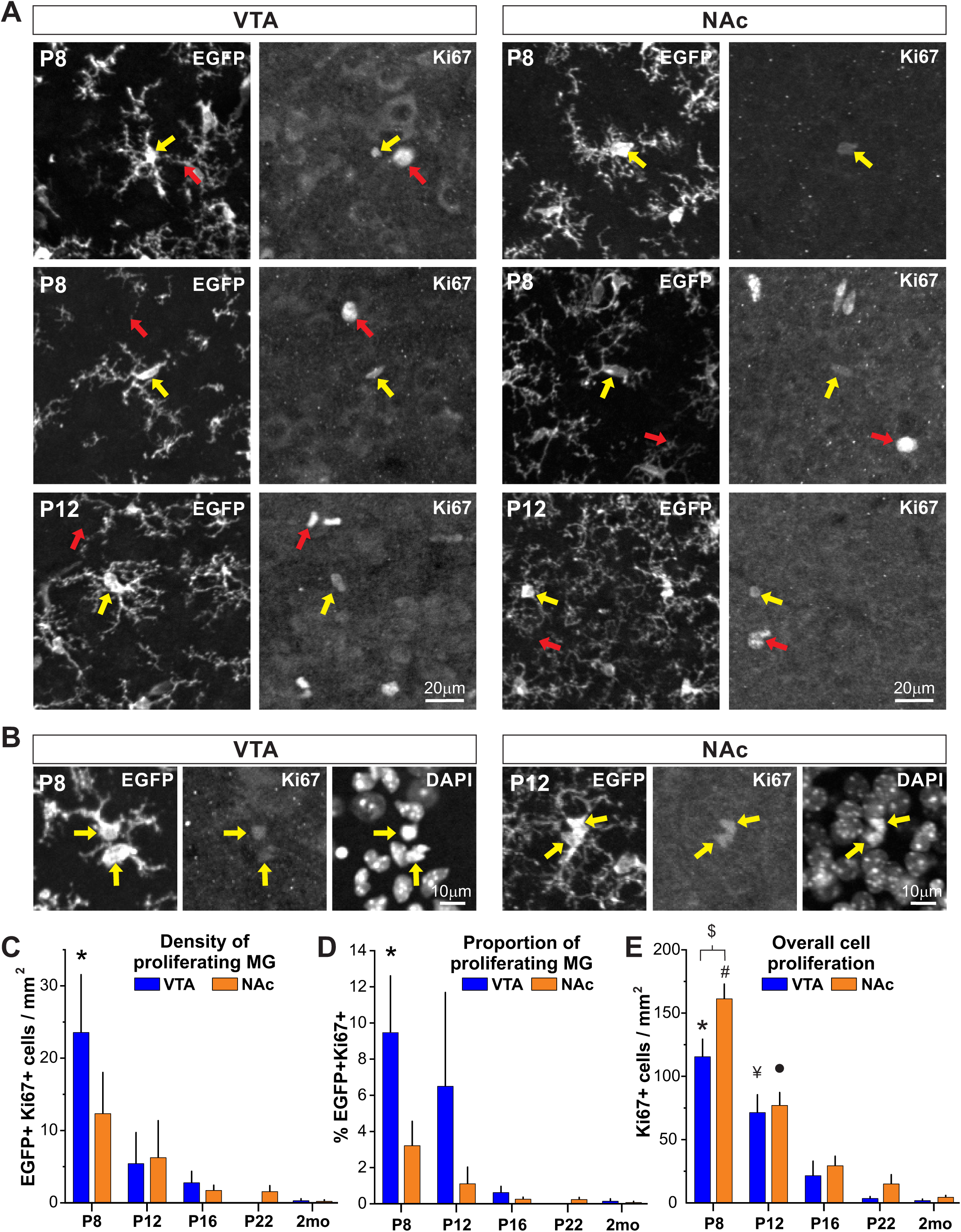
Microglial proliferation peaks during the first 2 postnatal weeks in the mesolimbic circuitry. **(A)** Representative images of immunohistochemical staining for Ki67, a marker of cell proliferation, and EGFP+ microglia at P8 and P12 in the VTA and NAc. *Yellow arrows* highlight examples of EGFP+Ki67+ cells and *red arrows* highlight examples of EGFP-Ki67+ cells. **(B)** Examples of EGFP+Ki67+ microglia in the VTA and NAc showing both morphological profiles of cell division and distinct DAPI-labeled nuclei. **(C)** Density of proliferating microglia n the VTA and NAc across early postnatal development. 2-way ANOVA; main effect of age, F(_4,42_) = 8.5, P = 0.00004; main effect of brain region, F(_1,42_) = 0.7, P = 0.4; age x brain region interaction, F(_5,42_) = 1.1, P = 0.4. P < 0.008 P8 VTA vs. P16 VTA, P8 VTA vs. P22 VTA, P8 VTA vs. 2mo VTA posthoc comparison. **(D)** Proportion of microglia that are proliferating at a given developmental time point in VTA and NAc. 2-way ANOVA; main effect of age, F(_4,42_) = 6.0, P = 0.0007; main effect of brain region, F(_1,42_) = 5.1, P = 0.03; age x brain region interaction, F(_5,42_) = 1.8, P = 0.14. * P < 0.02 P8 VTA vs. P16 VTA, P8 VTA vs. P22 VTA, P8 VTA vs. 2mo VTA posthoc comparison. **(E)** Density of all proliferating (EGFP+ and EGFP-) cells in the VTA and NAc across postnatal development. 2-way ANOVA; main effect of age, F(_4,42_) = 6.0, P < 0.00001; main effect of brain region, F(_1,42_) = 6.2, P = 0.02; age x brain region interaction, F(_5,42_) = 1.9, P = 0.13. * P < 0.00001 P8 VTA vs. P16 VTA, P8 VTA vs. P22 VTA, P8 VTA vs. 2mo VTA; # P < 0.00001 P8 NAc vs. P12 NAc, P8 NAc vs. P16 NAc, P8 NAc vs. P22 NAc, P8 NAc vs. 2mo NAc; ¥ P < 0.03 P12 VTA vs. P16 VTA, P12 VTA vs. P22 VTA, P12 VTA vs. 2mo VTA; • P < 0.046 P12 NAc vs. P16 NAc, P12 NAc vs. P22 NAc, P12 NAc vs. 2mo NAc; $ P8 VTA vs. P8 NAc posthoc comparisons.

Although proliferative microglia were abundant at P8 and P12, non-microglial proliferating cells were roughly 4 times more abundant (**Fig. 3E**). Similar to the time course of microglial proliferation, non-microglial cell proliferation was highest at P8 but did not fully decline until after P22. In contrast to proliferative dynamics within the microglial cell population, non-microglial proliferation declined more slowly in the NAc, suggesting that independent factors regulate microglial proliferation and proliferation of other cell populations. Consistent with this idea, there was no correlation between the abundance of proliferating microglia and the abundance of non-microglial proliferating cells in either the VTA or the NAc (**Fig. S3 C**). Occasionally we observed Ki67+ cells that appeared to be perivascular macrophages, which lacked processes, showed dimmer EGFP, and were located near large blood vessels (**Fig. S4 A**), but these were clearly distinguishable from ramified parenchymal microglia and represent less than 5% of observed EGFP^+^Ki67^+^ cells. We also observed instances in which microglia showed clear morphological profiles of proliferation but were Ki67^-^ (**Fig. S4 B**), indicating that our quantification underestimates the number of proliferating microglia at these time points. Collectively, these data support the idea that developmental increases in mesolimbic microglia are driven by local proliferation of resident microglia.

### Tissue expansion cannot fully account for decline in microglial density

What accounts for the sharp decline in microglial numbers in the VTA and NAc between the third postnatal week and adulthood? Brain volume and brain weight continue to increase throughout the first 6 postnatal weeks in mice (Semple *et al.*, 2013; Nikodemova *et al.*, 2015). Although this is believed be most prominent in the cortex, tissue expansion during this growth could also shape microglial density in the VTA and NAc and contribute to density decreases between the third postnatal week and adulthood. To determine the extent of developmental tissue expansion in these brain regions, we quantified the area of the VTA and NAc at key developmental time points (**Fig. 4A**). The area of the VTA did not change greatly between P10 and P25 but did increase significantly between P25 and 2 months (**Fig. 4B**). In contrast, area of the NAc increased steadily between P10 and 2 months (**Fig. 4C**). These trends were largely conserved throughout the anterior-posterior extent of both nuclei (**Fig. S5**). Within the VTA, the increase in nucleus area from P25 to 2 months represents a roughly 11% increase in size. This would be expected to decrease microglial density by approximately 9.9%. Yet, microglial density decreases by 36% during the same interval. Similarly, in the NAc, nucleus area increases 20% from P25 to 2 months, which would lead to a 16.6% reduction in cell density. However, NAc microglial density decreases 44% during this interval. These data suggest that tissue expansion contributes to but cannot fully account for the decrease in VTA and NAc microglial abundance between the third postnatal week and adulthood.

**Figure 4.**
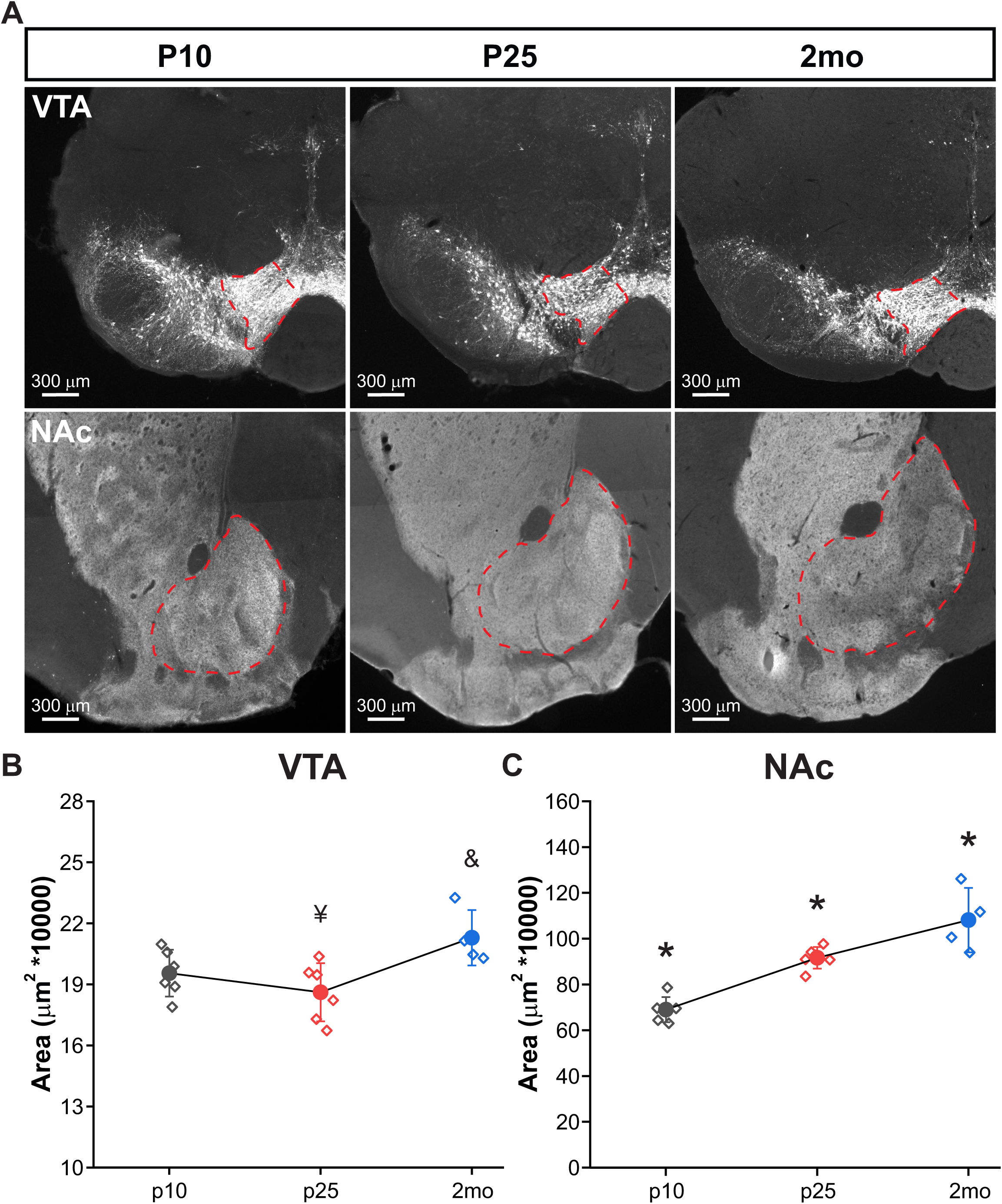
Moderate size expansion of VTA and NAc during development. **(A)** Representative images from coronal brain sections immunostained for tyrosine hydroxylase (TH), showing the size of the ventral tegmental area (VTA) and nucleus accumbens (NAc) (outlined with dotted *red lines*) at different developmental timepoints. **(B)** Quantification of VTA size during postnatal development, ANOVA F(_2,13_) = 5.02335, p=.02419. & P < 0.05 vs. P25, ¥ P < 0.05 vs. 2mo. **(C)** Quantification of NAc size during postnatal development, ANOVA F(_2,13_) = 29.3, p=1.54003E-5. * P < 0.05 vs all comparisons. Data points represent the average size of the brain region obtained for one mouse (N = 5-16 images analyzed per mouse and N=4-6 mice analyzed per time period).

### Mesolimbic microglia undergo programmed cell death to achieve mature densities

If tissue expansion cannot fully account for decreases in VTA and NAc microglial density, what mechanisms underlie this decline? FACS analysis of microglia in the whole brain suggested that cell death may accompany periods of microglial decline (Nikodemova *et al.*, 2015). To determine whether programmed cell death plays a role in decreasing microglial cell density between postnatal periods and adulthood, we examined caspase-3 staining in the VTA and NAc at key developmental time points. Although Casp3^+^ cells could occasionally be observed in the VTA and NAc at early time points, Casp3^+^ microglia were most abundant in the third postnatal week and beyond. Casp3^+^ microglia frequently showed morphological profiles indicative of programmed cell death, with prominent process blebbing (**Fig. 5A** and **Fig. S6A**). Although Casp3^+^ microglia were sometimes observed in close proximity to non-microglial Casp3^+^ cells (**Fig. 5B**), they did not cluster with other Casp3^+^ microglia and instead appeared scattered throughout the tissue. In addition, there was no obvious relationship between the location of Casp3^+^ microglia and local microglial density. For example, Casp3^+^ microglia did not appear to be located in regions that had a higher local abundance of microglial cells.

**Figure 5.**
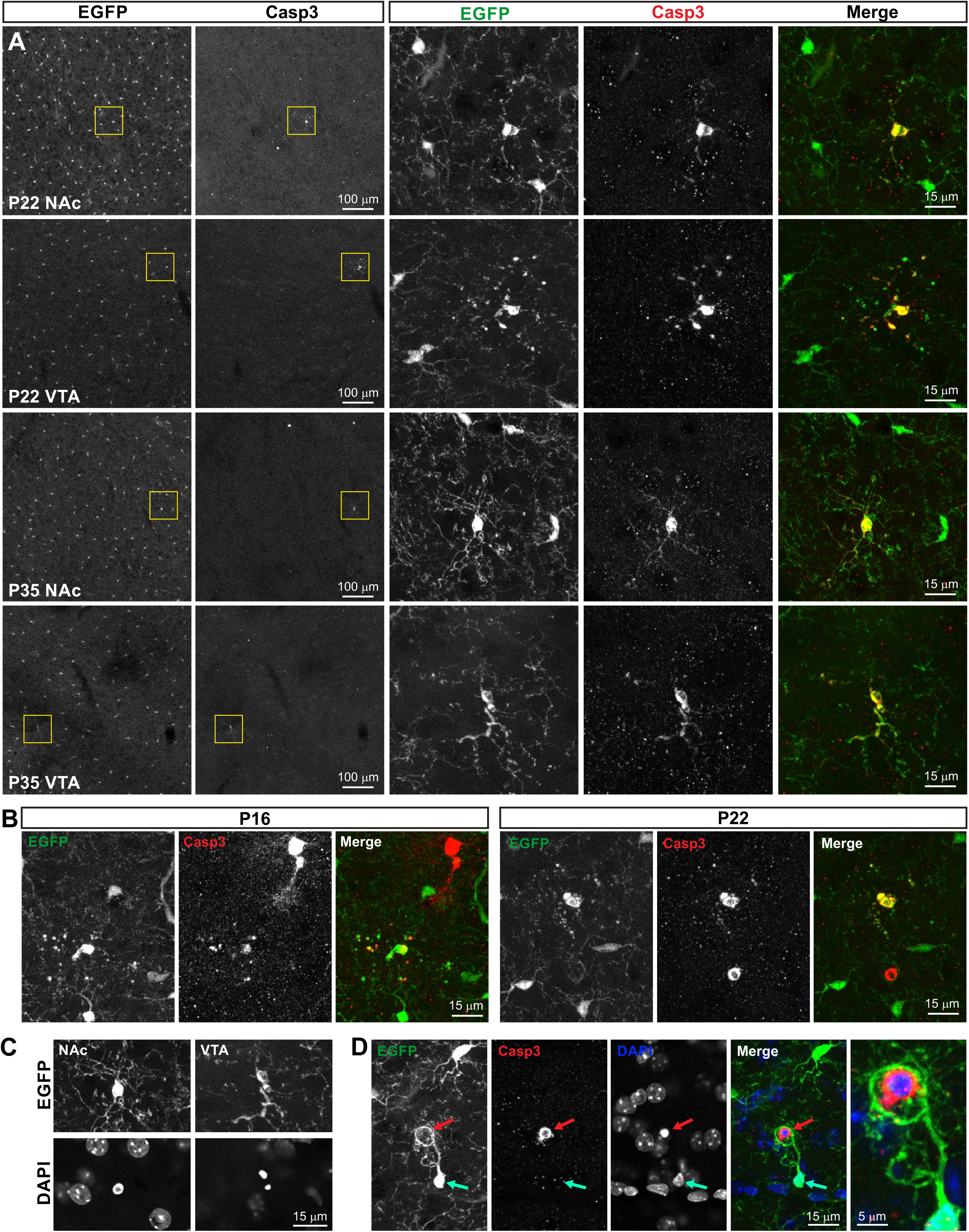
Mesolimbic microglia undergo programmed cell death during the 3^rd^ – 5^th^ postnatal weeks. **(A)** Representative images of immunostaining for caspase 3 (Casp3), a marker of cell death, and EGFP+ microglia in the VTA and NAc during the 3^rd^ and 5^th^ postnatal weeks. *Yellow boxes* indicate regions shown at higher magnification at right. Some EGFP+Casp3+ microglia also exhibit prominent membrane blebbing. **(B)** Examples of EGFP+Casp3+ microglia found in close proximity to non-microglial EGFP-Casp3+ cells. **(C)** DAPI staining for P35 NAc and VTA cells from *A* illustrating how EGFP+Casp3+ microglia also exhibit condensed chromatin typical for cells undergoing programmed cell death. **(D)** EGFP+Casp3+ microglia can clearly be distinguished from instances in which healthy EGFP+ microglial cells are engaging in phagocytotic engulfment of a neighboring Casp3+ cell. *Red arrow* highlights the Casp3+ nucleus and condensed chromatin of a dying cell being engulfed by an EGFP+ microglial phagocytotic cup. *Cyan arrow* highlights the absence of Casp3 labeling and the intact nucleus of the EGFP+ microglial cell engulfing the Casp3+ cell. Example shown is from P35 NAc. Not all z-levels for this max stack are shown in the DAPI channel for clarity.

Casp3^+^ dying microglia could be clearly distinguished from healthy microglia that were phagocytosing nearby non-microglial Casp3^+^ cells (**Fig. 5C** and **Fig. S6 B,C**). In these cases, phagocytosing microglia did not exhibit any membrane blebbing or chromatin condensation and instead formed a prominent phagocytotic cup around a nearby Casp3^+^ cell with condensed nuclei. In cases of microglial programmed cell death, Casp3+ microglia themselves exhibited condensed nuclei (**Fig. 5D** and **Fig. S6A**). Curiously, neighboring “healthy” microglia were never observed to phagocytose Casp3^+^ microglia, raising the question of how they are ultimately cleared from the tissue and suggesting that microglia in the developing brain exhibit specificity in what types of cellular debris they will clear from the surrounding tissue.

### Microglial functional interactions during periods of elevated density

Elevated microglial density during development could result from exuberant proliferation that temporarily outstrips regulatory mechanisms that establish proper microglial numbers in the adult VTA and NAc. Alternatively, microglia may be overproduced to carry out specific functional roles during discrete developmental windows. To gain initial insight into which of these possibilities is more likely, we quantified the prevalence of microglial phagocytotic cups and lysosomes in the developing VTA and NAc.

Microglia can shape maturing circuitry through elaboration of phagocytotic cups and engulfment of live, dying, or newborn cells (Brown & Neher, 2014; VanRyzin *et al.*, 2019). In the VTA and NAc microglial phagocytotic cups were easily recognized as large, 3-6um circular compartments within microglial processes and occasionally somas (**Fig. 6A,B**). Phagocytotic cups exhibited a rim of CD68+ lysosomes and in most instances contained condensed chromatin within the core of the cup, indicating engulfment of a dying cell (**Fig. 6C**). Microglia that possessed phagocytotic cups seemed to occur in spatially grouped clusters, rather than being evenly spread throughout the tissue and frequently a particular microglial cell displayed 2-3 phagocytotic cups, while neighboring cells had none. Microglia with phagocytotic cups were evident in the VTA and NAc of 100% of analyzed mice at P8 and P12, with the percentage falling steadily thereafter (**Fig. 6D**). More extensive quantification at P8 and P12 when these structures were most prevalent revealed a similar abundance of phagocytotic cups at both the tissue level and cellular level within the VTA and NAc (**Fig. 6E-G**). Collectively, these data indicate that microglia engulfment of dying cells peaks prior to the period of elevated microglial cell density. Further, the enhanced microglial overproduction observed in NAc is not correlated with more extensive engulfment of dying cells.

**Figure 6.**
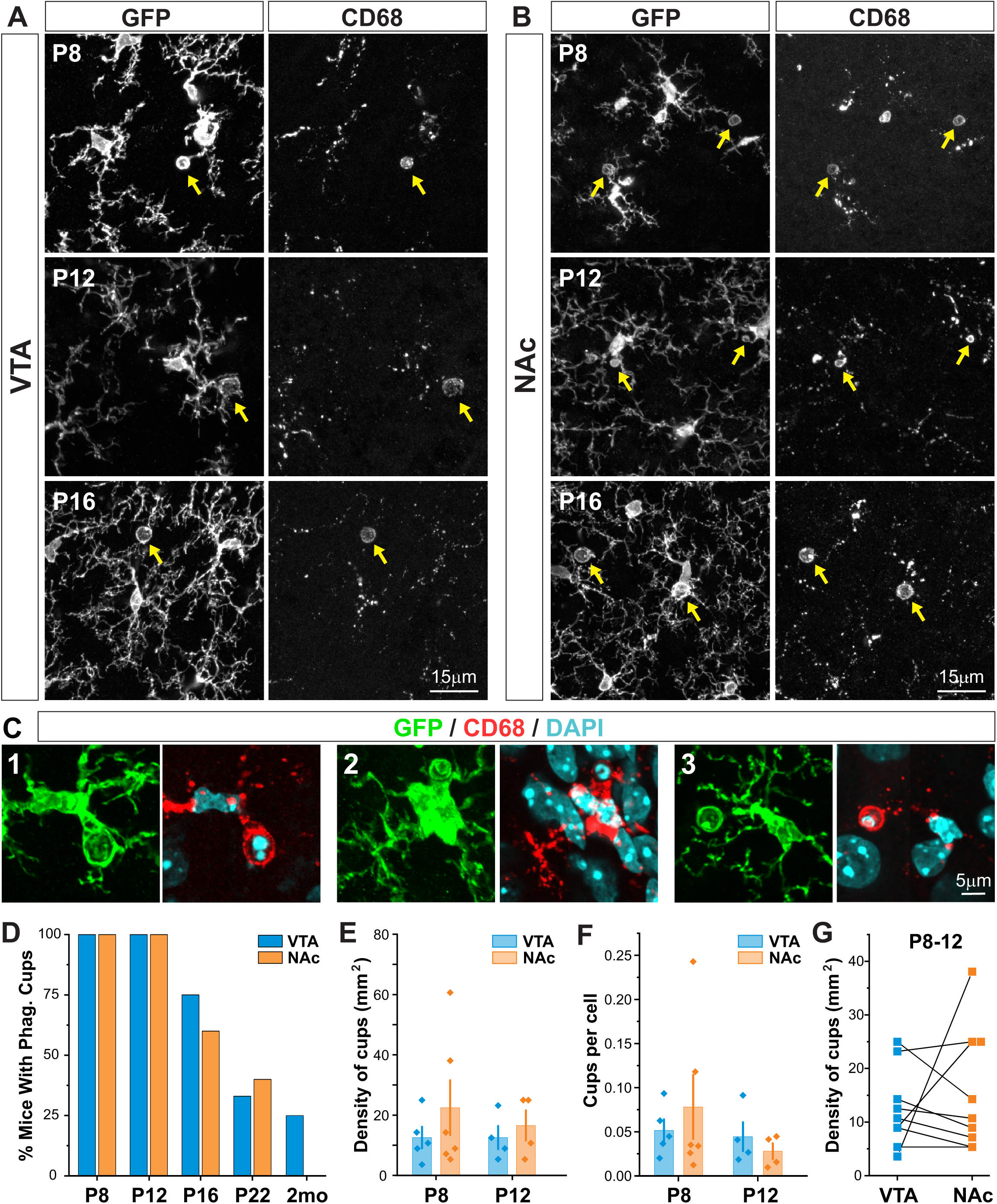
Microglial engulfment of dying cells in the mesolimbic circuitry peaks during the first two postnatal weeks. **(A,B)** Representative images from different developmental timepoints of VTA and NAc microglia that exhibit phagocytotic cups. *Yellow arrows* highlight the CD68 lysosome rim present in these structures. **(C)** Staining with DAPI shows condensed chromatin within phagocytotic cups indicating that these structures are formed around dying cells. **(D)** Percentage of mice at each developmental point that have microglia with phagocytotic cups. **(E)** Density of microglial phagocytotic cups in the VTA and NAc during the first two postnatal weeks. 2-way ANOVA; main effect of age, F(_1,15_) = 0.2, P 0.7; main effect of brain region, F(_1,15_) = 1.0, P = 0.3; age x brain region interaction, F(_2,15_) = 0.2, P = 0.7. **(F)** Quantification of phagocytotic cups per microglial cell in the VTA and NAc during the first two postnatal weeks. 2-way ANOVA; main effect of age, F(_1,15_) = 1.2, P 0.3; main effect of brain region, F(_1,15_) = 0.04, P = 0.8; age x brain region interaction, F(_2,15_) = 0.7, P = 0.4. Data points represent average values obtained for each mouse (N = 2-3 images analyzed per region per mouse and N=4-6 mice analyzed per time period). **(G)** Comparison of abundance of phagocytotic cups across brain regions within the same mouse. Paired T-test, P = 0.6.

In developing sensory systems, microglia phagocytose and/or trogocytose synaptic elements to support circuit maturation (Frost & Schafer, 2016; Weinhard *et al.*, 2018). This largely occurs during the first postnatal week and the abundance of microglial lysosomes correlates with peak synapse engulfment (Schafer *et al.*, 2012). At P8, microglial lysosome content in both VTA and NAc exceeded that observed in adult animals by roughly 2-3 fold (**Fig. 7A,B** and **Fig. S7**). In contrast to primary sensory circuits where microglial lysosome content drops after the first postnatal week, VTA and NAc microglial lysosome content remained elevated at P12 and P16, before declining nearly to adult levels by P22 (**Fig. 7B**). Although the time course of declining microglial lysosome content was similar across VTA and NAc, the levels of lysosome content were consistently higher in the NAc compared to the VTA; two-way ANOVA revealed significant main effects of age (F(_4,38_) = 32.1, P <0.00001) and brain region (F(_1,38_) = 6.2, P = 0.02), but no significant interaction between the two (F(_5,38_) = 0.2, P = 0.9). Quantification of tissue levels of CD68 also revealed regional differences. Because tissue levels of CD68 are determined both by lysosome content of individual cells and tissue density of cells, tissue content of CD68 in NAc significantly exceeded that of VTA (two-way ANOVA main effect of age F(_4,38_) = 26.2, P <0.00001; main effect of brain region F(_1,38_) = 4.7, P <0.00001; interaction F(_5,38_) = 3.2, P = 0.03), and showed a distinct peak at P16 (P16 NAc vs all other ages, P < 0.01), identifying a developmental window during which the NAc is particularly enriched in microglial lysosomes. Together these observations raise the possibility that refinement of developing circuitry is more extensive within the NAc and requires enhanced microglial overproduction during the second and third postnatal weeks compared to other brain regions.

**Figure 7.**
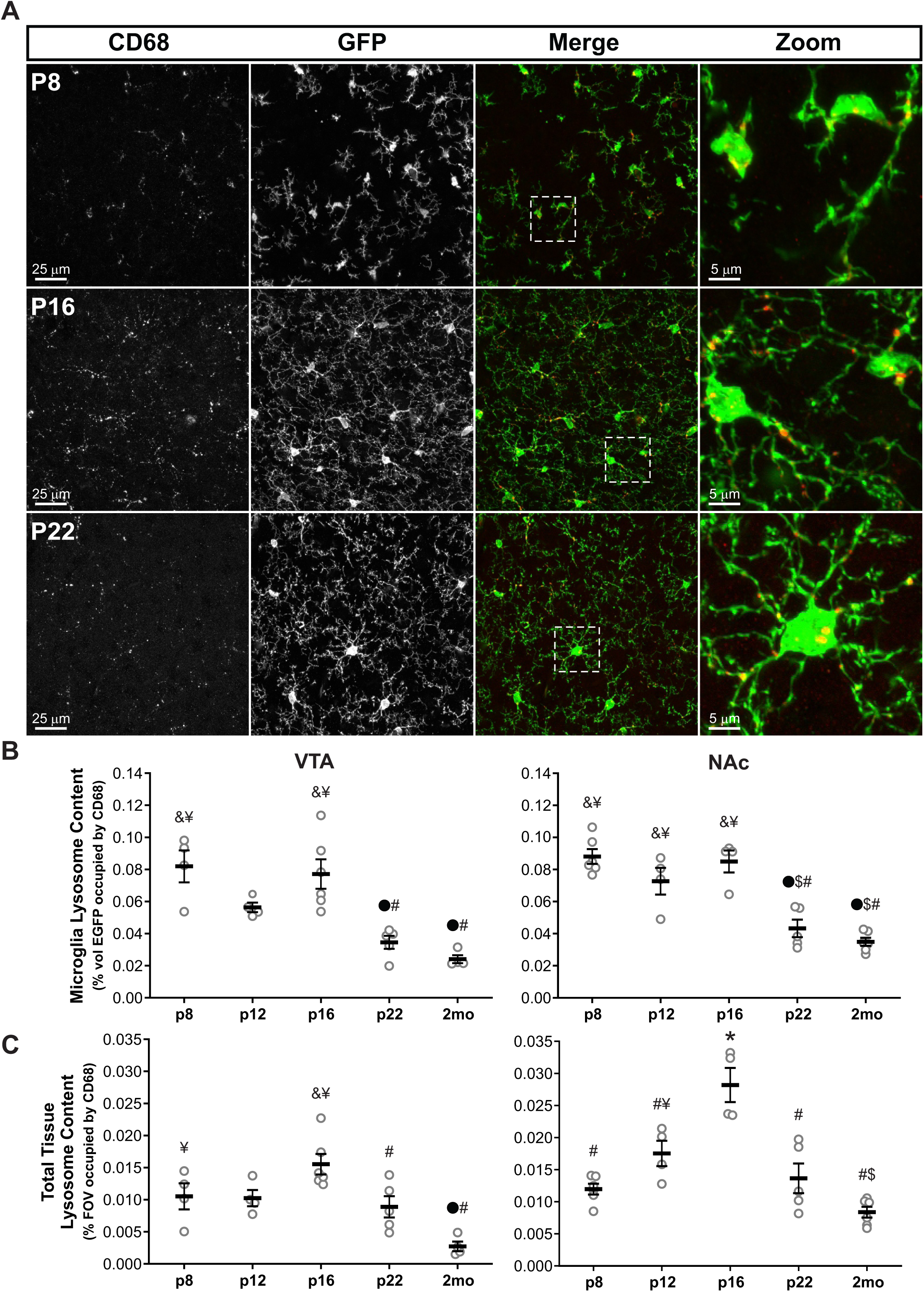
The developing NAc shows a discrete window of elevated microglial lysosome abundance. **(A)** Representative images from NAc of *CX3CR1*^*EGFP/+*^ mice immunostained for microglial specific lysosome membrane protein CD68. *Dashed white box* highlights region shown at higher magnification at right, illustrating the colocalization of CD68 and EGFP+ microglia. **(B)** Quantification of VTA and NAc microglial lysosome content (% volume of EGFP occupied by CD68) during early postnatal development. VTA ANOVA F(_4,18_) = 12.5, p = 0.00005. • P < 0.005 vs. P8, # P < 0.005 vs. P16, & P < 0.005 vs. P22, ¥ P < 0.001 vs. 2mo. NAc ANOVA F(_4,20_) = 22.6, p < 0.00001. • P < 0.001 vs. P8, $ P < 0.05 vs. P12, # P < 0.001 vs. P16, & P < 0.05 vs. P22, ¥ P < 0.001 vs. 2mo. **(C)** Quantification of total tissue content of microglial lysosomes (% of field of view occupied by CD68) during early postnatal development in the VTA and NAc. VTA ANOVA F(_4,18_) = 8.7, P = 0.0004. • P < 0.05 vs. P8, # P < 0.001 vs. P16, & P < 0.05 vs. P22, ¥ P < 0.05 vs. 2mo; NAc ANOVA F(_4,20_) = 18.6, P < 0.00001. * P < 0.01 vs. all comparisons, $ P < 0.05 vs. P12, # P < 0.01 vs. P16, ¥ P < 0.05 vs. 2mo. Data points represent average microglial lysosome content or tissue content of microglial lysosomes obtained for one mouse; N=2-3 images analyzed per mouse and 4-6 mice analyzed per time period.

## DISCUSSION

### Microglial colonization of the developing mesolimbic circuitry

Between the first and third postnatal weeks in mice, microglia transition from clustering in key hot spots to a tiled, ubiquitous distribution throughout the CNS (Squarzoni *et al.*, 2014; Reemst *et al.*, 2016). The time course of this phase of microglial maturation has not been fully mapped out. In previous studies, we found that microglia are evenly spread throughout basal ganglia nuclei by the end of the first postnatal week but at cell densities well below those observed in the adult (De Biase *et al.*, 2017). Six days later, however, they had attained densities that significantly exceeded those observed in the adult. Here, we extend these observations and find that VTA and NAc microglial cell density at P8 already significantly exceeds that observed at P6 and peak densities are attained during the second and third postnatal weeks in the NAc and VTA respectively. The sharp increase in microglial density at P8 suggests that expansion of the microglial population in these nuclei is not gradual but commences with a surge between P6 and P8 and ongoing expansion into the third postnatal week. Ki67 labeling and morphological profiles of dividing microglia indicate that this surge in microglial numbers is accomplished largely through cell proliferation. The high mouse to mouse variability observed in microglial Ki67 labeling and cell density at P8 support the idea that a proliferative surge occurs during a discrete developmental window; mice caught in the midst of this surge exhibited many Ki67+ microglia, while mice having completed or not yet initiated this surge exhibited few Ki67+ microglia.

What triggers this burst of microglial proliferation, particularly the dramatic increase of microglia in the NAc? We did not observe any correlation between the number of Ki67+ microglia and total Ki67+ cells in the vicinity, suggesting that independent factors regulate the proliferation of microglia and other cell populations and ambient trophic factors that affect numerous progenitor populations are unlikely to trigger this microglial proliferative surge. We also did not find obvious correlations between the number of Ki67+ microglia and local microglial density or spatial relationships between Ki67+ microglia and “gaps” in microglial tissue coverage (*data not shown*), suggesting that autocrine signaling within the microglial cell population is unlikely to play a dominant role in regulating this surge of microglia. Ki67+ microglia also did not appear to be spatially clustered or move in an obvious “wave” from one area to another. True pup locomotion begins around P7-P8 (Branchi & Ricceri, 2002; Dehorter *et al.*, 2011; Brust *et al.*, 2015), a behavioral milestone that will critically engage and shape neuronal activity within these circuits. Key events in maturation of the membrane properties and synaptic contacts of striatal medium spiny neurons also begin to occur during this window (Kozorovitskiy *et al.*, 2015; Peixoto *et al.*, 2016; Krajeski *et al.*, 2018). A key area of future study will be to determine whether these changes in behavior and circuit activity are causally linked to maturation of microglia within these regions.

### Microglia undergo developmental overproduction and refinement analogous to other CNS cell populations

During development, neuroectoderm derived neurons, astrocytes, and oligodendrocyte lineage cells are produced in excess and a subset of these cells then undergo programmed cell death. This overproduction and refinement is believed to be a critical mechanism for optimizing CNS cellular composition, removing any defective cells, and ensuring correct matching of signaling partners in the mature CNS (Yamaguchi & Miura, 2015). Microglia arise from a distinct cell lineage, primitive macrophage progenitors from the yolk sac that invade the CNS during embryonic periods (Ginhoux *et al.*, 2010, 2013). Whether similar overproduction and refinement mechanisms are critical for determining the mature microglial population has not been directly demonstrated.

In the adult CNS, long term *in vivo* imaging and clonal analysis have shown that microglia proliferate and undergo cell death to maintain consistent numbers (Askew *et al.*, 2017; Füger *et al.*, 2017; Réu *et al.*, 2017; Tay *et al.*, 2017). Reports from several brain regions have also shown that microglial density increases during early postnatal development but then declines by adulthood (Kim *et al.*, 2015; Askew *et al.*, 2017; Hagemeyer *et al.*, 2017; Riquier & Sollars, 2017). However, whether this decrease in microglial abundance was due to tissue growth or other mechanisms was not investigated. FACS sorting approaches also found increases and subsequent decreases in whole brain microglial numbers (Nikodemova *et al.*, 2015) and staining with DAPI and ssDNA suggested that microglial cell death may contribute to these decreases in cell number. However, cell counts and cell death in FACS protocols can also be influenced by variation in tissue processing or relative susceptibility of cells to death during processing at different ages.

We found that microglial density peaks within the mesolimbic system during the first 3 weeks of postnatal development and then declines to adult levels. Quantification of the size of the VTA and NAc throughout postnatal development indicated that the majority of tissue growth occurs prior to decreases in microglial cell numbers. Nonetheless, roughly 1/3 of the observed decrease in microglial cell density could be accounted for by tissue growth. This analysis indicated that other mechanisms must contribute to decline of microglial cell numbers between the third postnatal week and adulthood. Morphological analysis, assessment of chromatin condensation, and immunostaining with caspase-3 provided direct confirmation that microglial cell death contributes to this reduction in microglial density. Thus, microglia, like neuroectoderm-derived cell populations in the CNS, are overproduced and refined to attain mature cell population composition.

What triggers microglial cell death during development? In the adult, microglial proliferation and microglial cell death are spatially and temporally coupled to maintain consistent adult densities (Askew *et al.*, 2017), suggesting an equilibrium driven by availability of key trophic factors and/or autoregulatory signaling within the microglial cell population. Consistent with this, following microglial ablation in the adult CNS, repopulating microglia overshoot microglial numbers seen in control mice (Huang *et al.*, 2018) and excess microglia are removed through cell death to reestablish target microglial density (Zhan *et al.*, 2019). In the developing VTA and NAc, microglial proliferation and cell death are largely sequential and separated in time and Casp3+ microglia were not observed near to morphological profiles of proliferating microglia. However, Casp3+ microglia were distributed throughout the tissue, suggesting that, after the initial burst of microglial proliferation, evolving availability of ambient trophic factors could still play a critical role in determining the number of microglia that survive into adulthood. The identity of such trophic factors and what determines target densities of microglia in distinct adult brain regions are critical areas for ongoing study (De Biase & Bonci, 2018).

Curiously, neighboring microglia were never observed to engulf nearby dying microglia, raising the question of how these cells are ultimately cleared from the tissue. Similar observations have been made following microglial ablation in adult mice; dying microglia can clearly be observed (De Biase *et al.*, 2017; Zhan *et al.*, 2019), but how these cells are cleared has not been demonstrated. In the developing VTA and NAc, we readily observed many examples of microglia engulfing non-microglial cells undergoing programmed cell death. We also found examples of non-microglial Casp3+ cells that were not being engulfed by microglia. These observations indicate that microglia do not indiscriminately engulf any dying cells in their vicinity, but, rather, show specificity for which cell types they clear from the tissue.

### Maturation of other microglial features relative to cell numbers

Between the first and third postnatal weeks, microglia undergo massive changes in gene expression (Butovsky *et al.*, 2014; Bennett *et al.*, 2016; Matcovitch-Natan *et al.*, 2016; Hammond *et al.*, 2019; Li *et al.*, 2019), highlighting the magnitude of phenotypic changes in this cell population during this period of maturation. In agreement with the idea of broad phenotypic changes in microglia during this developmental window, we found that changes in microglial density were accompanied by significant changes in microglial morphology and lysosome content throughout the first three postnatal weeks. Morphologically, VTA and NAc microglia transitioned from amoeboid or sparsely branched cell structure at early postnatal periods to highly-ramified morphologies by the third postnatal week, with corresponding increases in microglial tissue coverage. Lysosome content in VTA and NAc microglia remained high throughout the first two postnatal weeks and then began to decline to adult levels by the third postnatal week. Although cell density, cell morphology, and lysosome content all underwent significant changes during the first three postnatal weeks, the precise time course of changes in each cellular attribute were not perfectly aligned. For example, morphological maturation generally lagged behind the rapid developmental increases in cell density during the second postnatal week. Further, lysosome content began to decline in the third postnatal week, prior to decreases in cell density. Together these observations reveal dynamic changes in numerous microglial attributes during this period and suggest that maturation of distinct microglial features is independently regulated. These observations also suggest that the progression toward adult microglial phenotypes may involve many sequential processes rather than one concerted switch from “early postnatal microglia” to “adult microglia.”

### Regional differences in the developmental maturation of microglia

In the adult CNS, microglia in the VTA and NAc display regionally specialized phenotypes, differing significantly in cell density, morphology, lysosome content, membrane properties, and transcriptome (De Biase *et al.*, 2017). Here, we show that these regional differences begin to emerge by P8 and that the time course and dynamics of microglial maturation also differ between these two regions. Microglia in the NAc undergo an explosive burst of proliferation in the second postnatal week and achieve a more significant overproduction relative to adult NAc microglia. In contrast, microglial proliferation and density increases in the VTA are more gradual and the magnitude of microglial overproduction less pronounced. Microglial overproduction in the NAc also appears to exceed that reported for microglia in other regions such as the brainstem (Riquier & Sollars, 2017) and cortex (Askew *et al.*, 2017; Hagemeyer *et al.*, 2017) and be comparable to that observed in hippocampus (Kim *et al.*, 2015). Cerebellum is the only region where reported microglial overproduction exceeds the levels we find in the NAc (Hagemeyer *et al.*, 2017). The time course of decline in microglial cell numbers may also vary significantly across regions; most other regions report significant decreases in microglial cell numbers by P21 (Kim *et al.*, 2015; Nikodemova *et al.*, 2015; Askew *et al.*, 2017; Hagemeyer *et al.*, 2017; Riquier & Sollars, 2017), whereas microglial density in the VTA and NAc does not decline significantly until after this time. Regional differences in morphological maturation are also evident. Both VTA and NAc microglia transition from simplified to more complex, ramified morphologies. However, increases in branching complexity and tissue coverage are more rapid in the NAc, and cell processes are over produced and refined to adult levels, a reduction that is not observed in the VTA. Collectively, these observations indicate that microglia in different brain regions follow distinct developmental trajectories as they move toward establishing the regionally specialized phenotypes observed in the adult.

What mechanisms give rise to these distinct developmental trajectories? Given the potency of local CNS cues in programing phenotypes of adult microglia (De Biase *et al.*, 2017; Bennett *et al.*, 2018), it seems likely that regional differences in local cues contribute to distinct maturational trajectories of these cells during the first three postnatal weeks. Relative to adult microglia, both neonatal microglia and adult repopulating microglia express low levels of surface molecules that may mediate contact inhibition (Zhan *et al.*, 2019). Regional differences in upregulation of these molecules could also shape regional variability in attainment of mature microglial cell numbers and branching patterns.

### Functional roles of microglia in the developing mesolimbic system

Microglia have been proposed to carry out numerous functional roles during CNS development and perturbation of microglial maturation is implicated in neurodevelopmental disorders (Frost & Schafer, 2016; Reemst *et al.*, 2016; Mosser *et al.*, 2017; Tay *et al.*, 2018). However, functional roles for microglia in the developing mesolimbic system have not been extensively investigated. Male/female differences in microglial functions during development have been reported for multiple brain regions (VanRyzin *et al.*, 2018, 2019), including the NAc during adolescence (Kopec *et al.*, 2018). We did not observe any obvious male female differences in the parameters we measured (**Fig. S8**), but this does not rule out that there are important male female differences in microglial features in these brain regions during these developmental windows.

In the neonatal cortex, microglia regulate survival of pyramidal neurons through release of insulin-like growth factor (IGF) (Ueno *et al.*, 2013). Brain-wide transcriptome analyses indicate that microglial expression of Igf1 remains elevated through P14 (Bennett *et al.*, 2016), suggesting that microglial release of this factor could influence neuronal survival in other brain regions. In addition, between P7 and P21, microglia express Igf2, Tgfb1, Vegfb, Cntf, Pdgfb, and Pdgfa (Bennett *et al.*, 2016) and microglia are a primary cellular source for many of these factors (Zhang *et al.*, 2014). Vegfb and Tgfb can shape dopamine neuron survival and dendrite outgrowth (Luo *et al.*, 2016; Caballero *et al.*, 2017) and both factors have been implicated in synaptogenesis and synaptic plasticity in multiple brain regions (Tillo *et al.*, 2012; Poon *et al.*, 2013). These observations, together with the elevated VTA and NAc microglial density during the second and third postnatal weeks, suggest a likely role for microglial secreted factors in shaping cell viability and circuit wiring in the developing mesolimbic system.

Microglia in the developing and adult CNS respond to a wide array of insults and perturbations. In some circumstances these microglial responses are neuroprotective, but in others they are detrimental or neurotoxic (Ransohoff & Perry, 2009; Sierra *et al.*, 2013). Additional research will be needed to clarify whether the pronounced elevation in microglial density in the NAc during the second and third postnatal weeks provides protection for surrounding neurons or instead represents a window of vulnerability during which insults are more detrimental. Striatal neurons are particularly susceptible to damage and death following hypoxia-ischemia (H/I) (Northington *et al.*, 2001; Mueller-Burke *et al.*, 2008). In these experiments, H/I is delivered around P7 and sequalae develop over the next week, suggesting that high microglial density in the second postnatal week could contribute to damage in this model. Koh and colleagues found that elevated developmental density of microglia in the hippocampus was correlated with increased susceptibility to seizures (Kim *et al.*, 2015), again suggesting that elevated developmental microglial density may be a liability rather than providing protection.

The highly ramified processes of adult microglia have been associated with a tissue surveillance phenotype (Madry & Attwell, 2015). Motile processes of adult microglia also appear to interact with synapses and can promote synapse formation, synapse elimination, and changes in synaptic receptor expression (Wu *et al.*, 2015). By the third postnatal week, we find that VTA and NAc microglia show morphologies highly comparable to that of adults. Whether tissue surveillance is less effective during early postnatal periods before microglia become fully ramified is unclear. Furthermore, the exact relationship between microglial process ramification and the degree to which microglia interact with and influence surrounding synapses has not been clarified. In the NAc, microglial process branching increases more steeply that in the VTA and tissue coverage briefly exceeds that observed in adults. This period of rapid increase in morphological complexity coincides closely with periods of synaptogenesis, spinogenesis, and maturation of synaptic activity within the striatum (Kuo & Liu, 2019). Synapse refinement instead, begins in the third postnatal week and may continue throughout adolescence (Kopec *et al.*, 2018; Kuo & Liu, 2019), suggesting that NAc microglial morphological maturation may be functionally aligned with synaptogenesis and spinogenesis.

Amoeboid morphology of microglia and elevated lysosome content in early postnatal development has been associated with phagocytotic activity of these cells in other brain regions (Frost & Schafer, 2016). Consistent with these findings, we observed microglia with phagocytotic cups containing cellular material, with cups being most prevalent in the first two postnatal weeks. Microglia in the developing amygdala phagocytose otherwise viable newborn cells that are destined to become astrocytes (VanRyzin *et al.*, 2019). Although we did not determine the identity of cells engulfed by microglia in the VTA and NAc, these are unlikely to be viable newborn cells as they exhibit condensed chromatin and stain positive for caspase 3, and we did not observe clear examples of microglial engulfment of Ki67+ cells. At P8, VTA and NAc microglial lysosome content was slightly lower than that reported for neonatal visual circuits (Schafer *et al.*, 2012), but remained elevated throughout the second postnatal week, suggesting that microglial involvement in circuit refinement may occur over a prolonged period in these brain regions. In the NAc, elevated microglial density combined with elevated cellular lysosome content to give rise to a unique peak in tissue CD68 at P16, highlighting this period as a key potential window for microglial involvement in circuit wiring.

### Conclusion

These studies provide foundational information about the maturation of microglia within the mesolimbic system and highlight that microglia in the VTA and NAc follow distinct developmental trajectories. These findings help identify potential regional distinctions in microglial contributions to circuit maturation and highlight temporal windows during which perturbation of microglia may have detrimental effects that could contribute to neurodevelopmental disorders involving these circuits.

## ACKNOWLEDGEMENTS

This work was supported by start-up funds from the David Geffen School of Medicine at UCLA (L.M.D.), the Brain and Behavior Research Foundation (L.M.D.), and the National Institute on Drug Abuse Intramural Research Program (A.B.). The authors thank the NIDA IRP Histology Core for tissue sectioning, A. Fernandez for contributions to experiments, and E. Moca for editorial suggestions during manuscript preparation.

## AUTHOR CONTRIBUTIONS

Conceptualization, L.M.D. and I.A.H.; Investigation, I.A.H., K.T.H, and L.M.D.; Formal Analysis, K.T.H., I.A.H., and L.M.D.; Visualization, K.T.H. and L.M.D.; Writing, L.M.D.; Funding Acquisition, L.M.D and A.B.; Supervision, L.M.D. and A.B.

## DATA ACCESSIBILITY

The original data files are available by contacting the last author.

## ABBREVIATIONS

NAc: Nucleus Accumbens
VTA: Ventral Tegmental Area
EGFP: Enhanced Green Fluorescent Protein
PFA: Paraformaldehyde
PBS: Phosphate Buffered Saline
NDS: Normal Donkey Serum
RT: Room Temperature
FACS: Fluorescence-Activated Cell Sorting

