## Supplementary Figures for "Mesolimbic microglia display regionally distinct developmental trajectories"

### Supplementary Figure 1, Related to Fig. 1

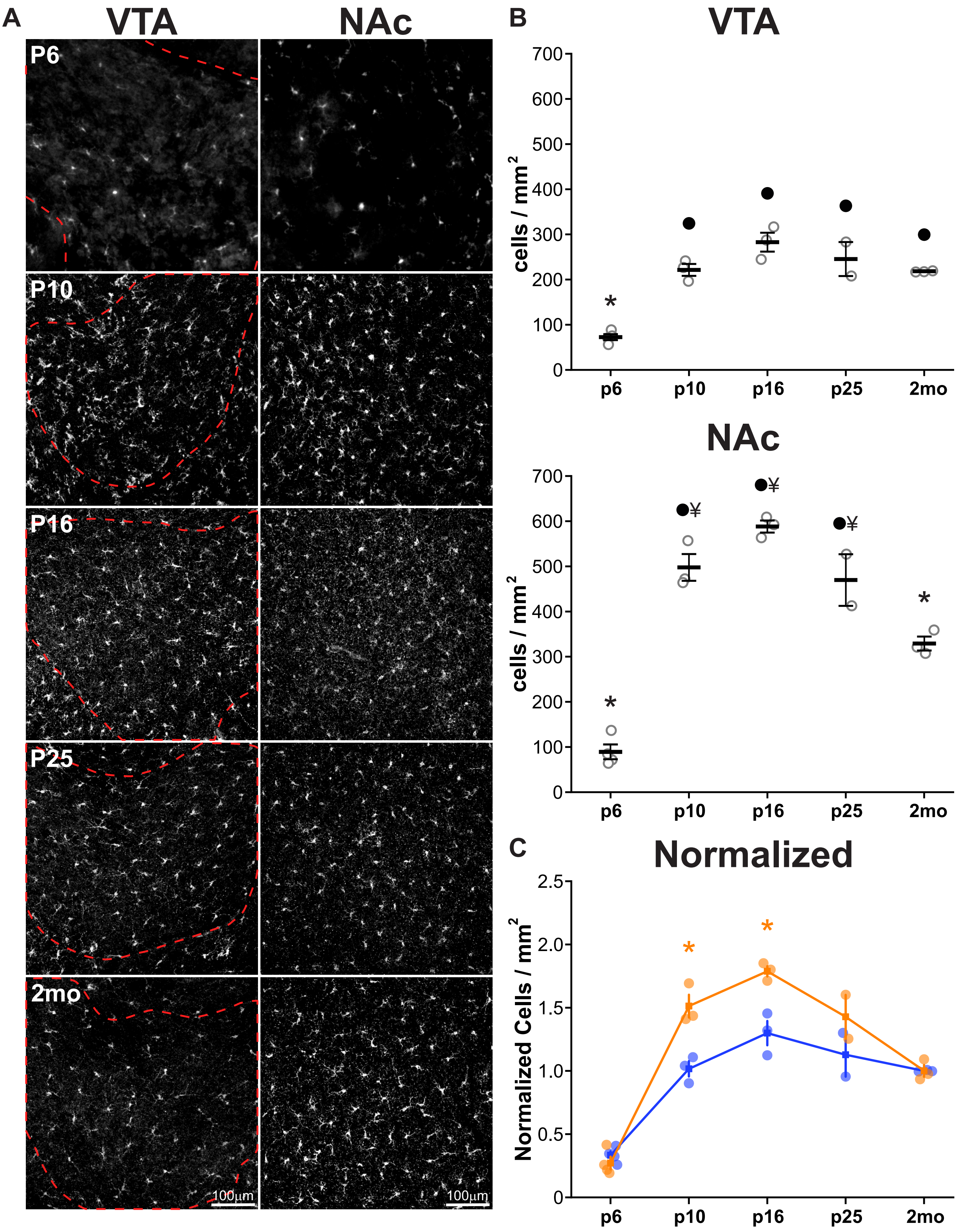

**Supplementary Figure 1. Density of mesolimbic microglia during development in wildtype mice. (A)** Representative images from the ventral tegmental area (VTA) and nucleus accumbens (NAc) of wildtype mice immunostained for ionized calcium binding adapter molecule 1 (Iba1), which is expressed specifically in microglia, at postnatal days (P) 6, 10, 16, 25, and 2mo. Boundary of VTA outlined by *dotted red line*. **(B)** Quantification of microglia density during early postnatal development in the VTA and NAc, with each data point representing the average microglial density for one mouse. N = 2-3 images per mouse, 2-4 mice per time point. VTA ANOVA $F_{(4,10)} = 32.5$, p=1.04534E-5. * P < 0.001 vs all comparisons, ● P < 0.001 vs. P6. NAc ANOVA $F_{(4,10)} = 75.8$, p=1.93523E-7. * P < 0.05 vs all comparisons, ● P < 0.001 vs. P6, ¥ P < 0.05 vs. 2mo. **(C)** Developmental microglial density normalized to the average value for each region at 2mo. * P < 0.005 P10 NAc vs VTA, P16 NAc vs VTA.

### Supplementary Figure 2, Related to Fig. 2

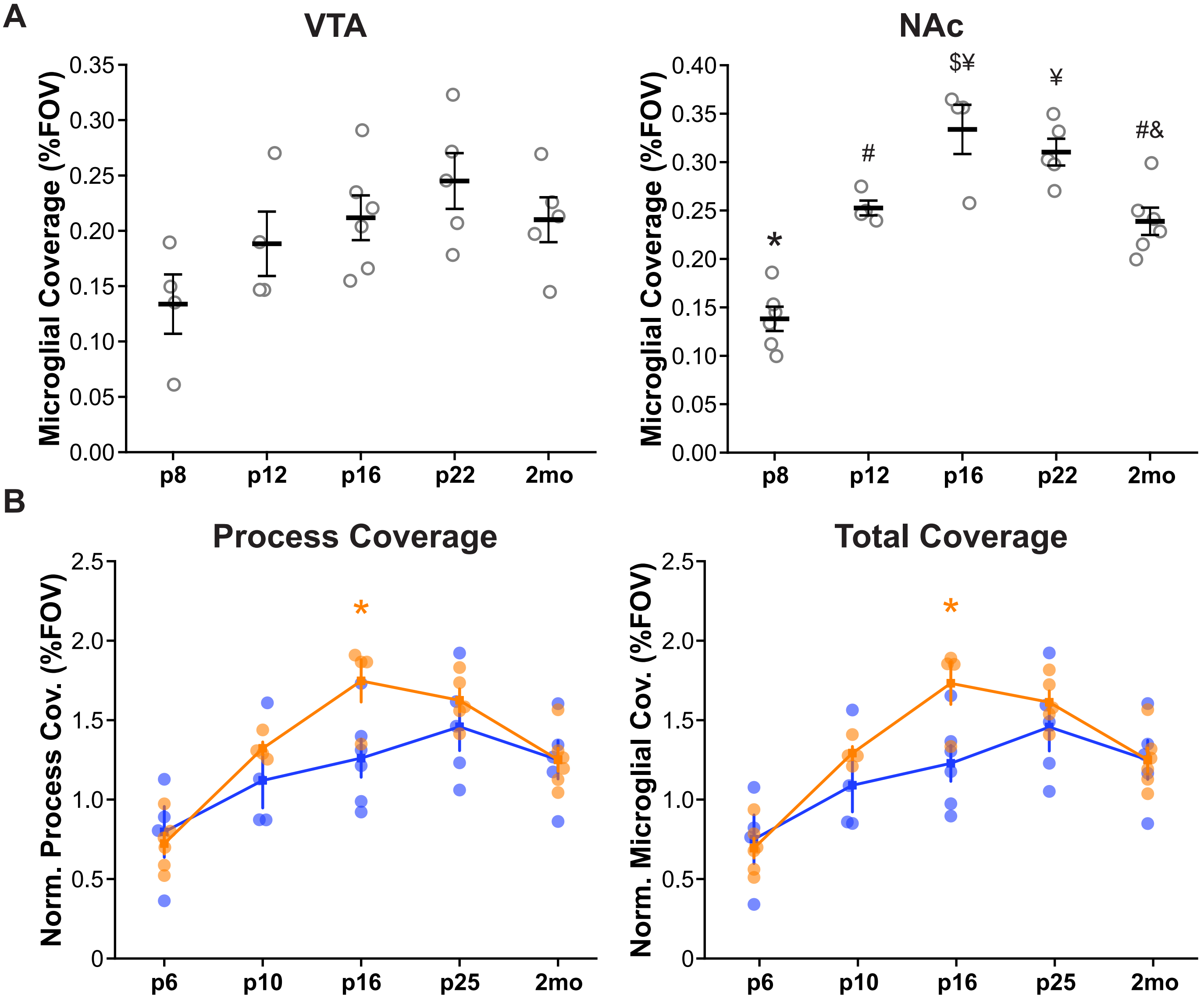

**Supplemental Figure 2. Total microglial tissue coverage of developing VTA and NAc.** Quantification of total microglial tissue coverage (cell processes + somas) within the VTA and NAc during early postnatal development; each data point represents average values obtained for one mouse. N=2-3 images per mouse, 4-6 mice per time point. VTA: ANOVA $F_{(4,19)} = 3.7$, p=.06332. NAc: ANOVA $F_{(4,20)} = 26.5$, p=9.69452E-8. * P < 0.001 vs. all comparisons, ● P < 0.05 vs. P8, $ P < 0.05 vs. P12, # P < 0.05 vs. P16, & P < 0.05 vs. P22, ¥ P < 0.05 vs. 2mo.

### Supplementary Figure 3, Related to Fig 3.

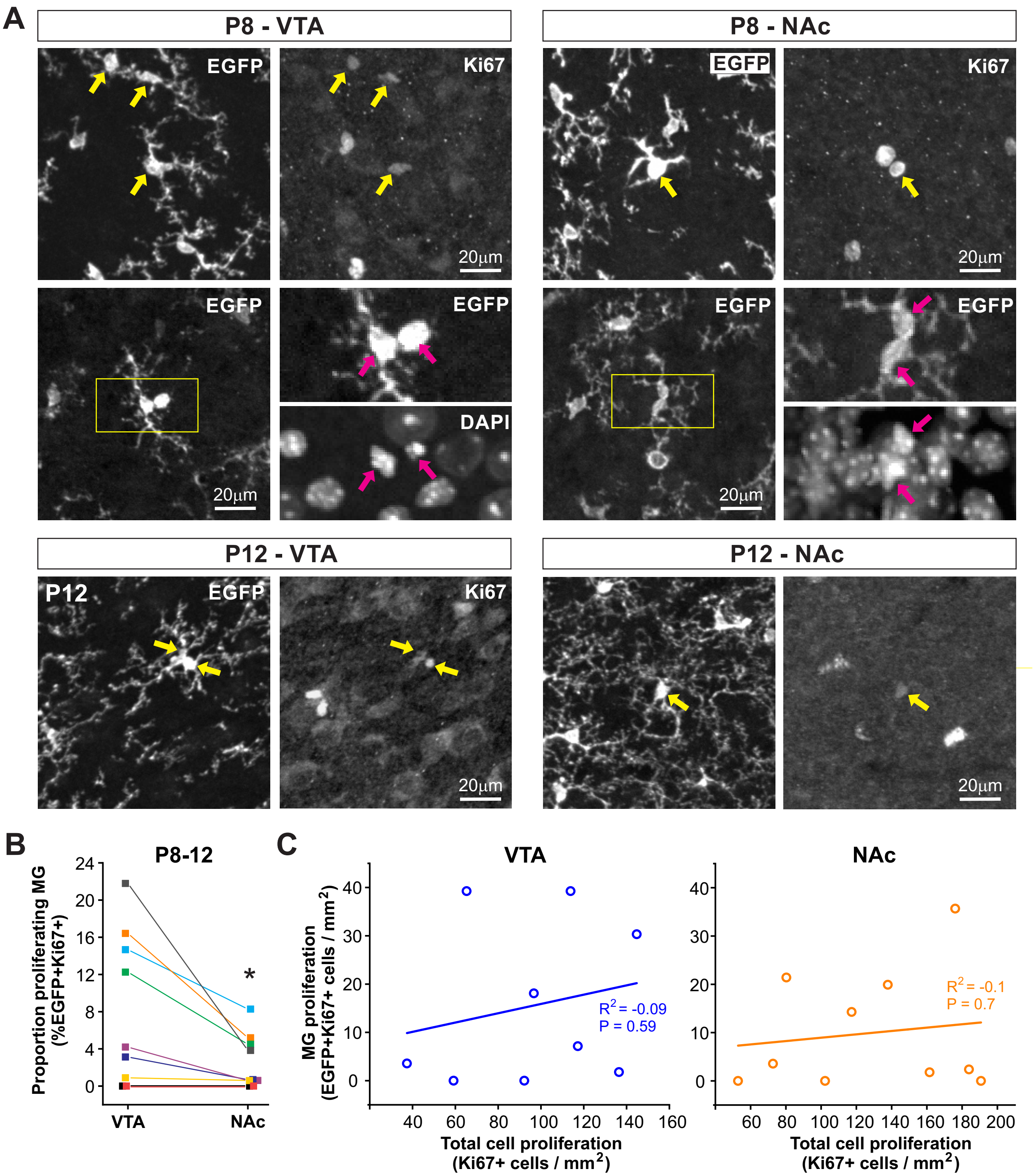

**Supplementary Figure 3. Microglial proliferation during postnatal development does not correlate with overall cell proliferation. (A)** Additional examples of Ki67+ microglia observed in the VTA and NAc during the first two postnatal weeks and cells that show morphological profiles indicative of cell division and distinct separate nuclei as visualized by DAPI staining. *Yellow arrows* indicate EGFP+ Ki67+ microglia. Region highlighted by *yellow box* shown at higher magnification at right. *Magenta arrows* indicate distinct nuclei of dividing cells. **(B)** Comparison of abundance of EGFP+Ki67+ microglia across brain regions within the same mouse. * $P = 0.03$, Paired T-test. **(C)** Relationship between microglial cell proliferation and overall cell proliferation in the VTA and NAc during early postnatal development. Data points represent average values for individual P8 and P12 mice.

### Supplementary Figure 4, Related to Fig. 3

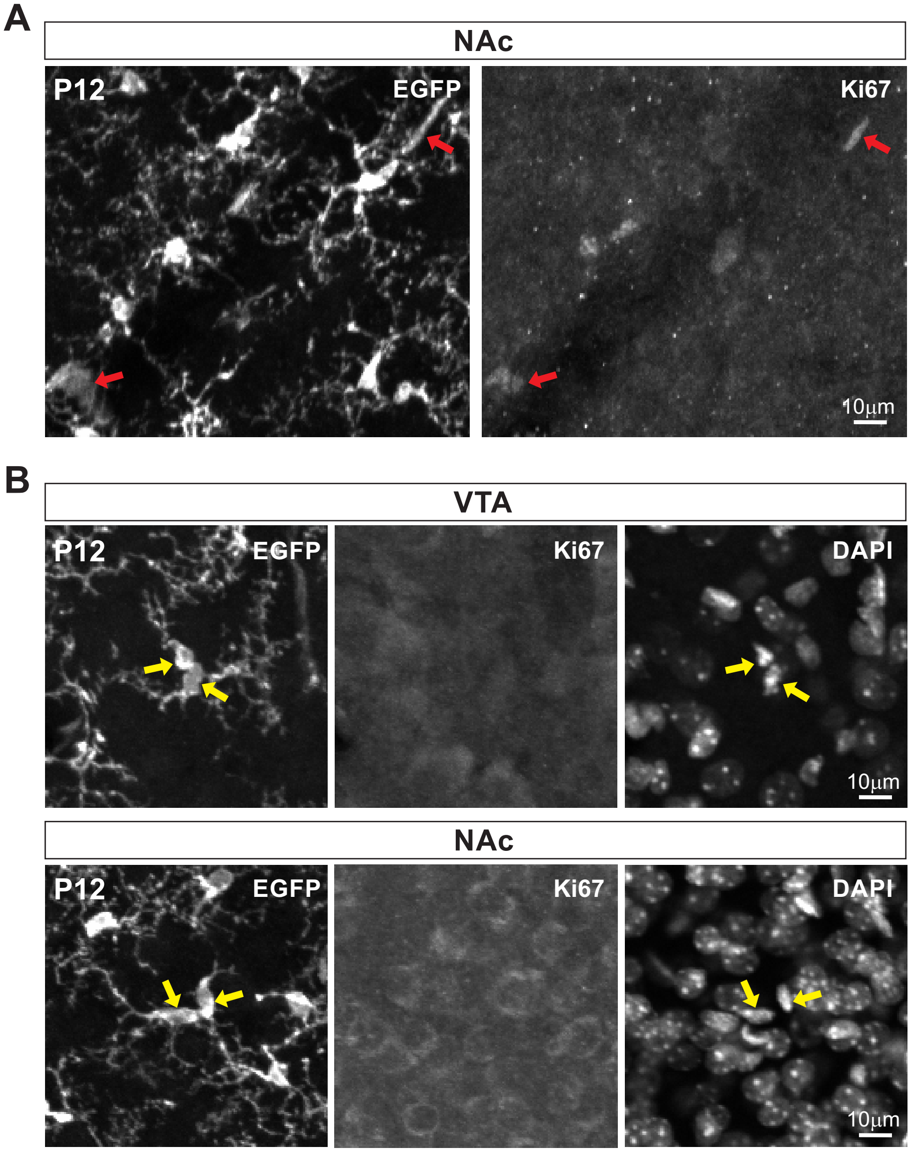

**Supplementary Figure 4. Ki67 staining reveals macrophage proliferation and does not label all proliferating microglia. (A)** Examples of EGFP+Ki67+ macrophages. These cells exhibited dimmer EGFP fluorescence, amoeboid morphology without any ramified processes and were generally found near major blood vessels and could thus easily be excluded from analysis. **(B)** EGFP+ microglia showing clear morphological profiles of cell division and distinct nuclei that do not stain positive for Ki67.

### Supplementary Figure 5, Related to Fig. 4

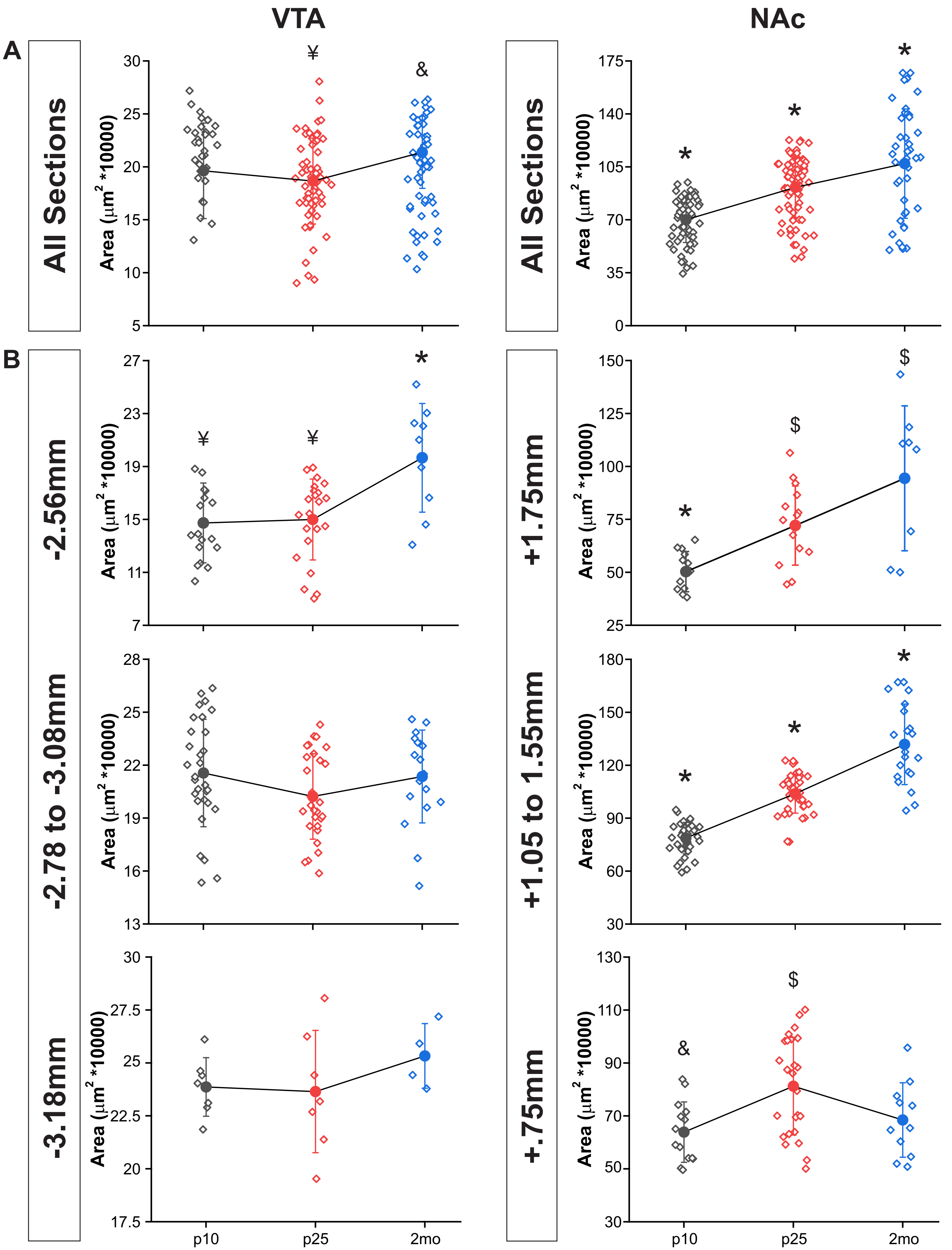

**Supplementary Figure 5. Growth of VTA and NAc nuclei during development. (A)** Quantification of VTA and NAc size during early postnatal development at postnatal days (P) 10, 25, and 2mo; each data point represents VTA or NAc size in individual brain sections throughout the rostral-caudal extent of the nucleus. N = 31-80 brain sections from 4-6 mice at each time period. VTA ANOVA $F_{(2,141)} = 4.5$, p=.01334. & $P < 0.05$ vs. P25, ¥ $P < 0.05$ vs. 2mo. NAc ANOVA $F_{(2,181)} = 33.9$, p=3.13881E-13. * $P < 0.005$ vs. all comparisons. **(B)** Same data points shown in *A* separated according to rostral-caudal location within the nucleus. Location relative to Bregma is listed for reference. **(B)** Rostral VTA ANOVA $F_{(2,45)} = 9.0$, p=5.19915E-4. * $P < 0.005$ vs. all comparisons, ¥ $P < 0.005$ vs. 2mo. Middle VTA ANOVA $F_{(2,74)} = 1.9$, p=.15112. Caudal VTA ANOVA $F_{(2,15)} = 0.8641$, p= 0.44138. Rostral NAc ANOVA $F_{(2,31)} = 10.7$, p=2.96041E-4. * $P < 0.05$ vs. all comparisons, \$ $P < 0.05$ vs. P10. Middle NAc ANOVA $F_{(2,97)} = 104.1$, p=7.08062E-25. * P < 1E-10 vs. all comparisons. Caudal NAc ANOVA $F_{(2,47)} = 6.1$, p=.00442. \$ $P < .005$ vs. P25, # $P < 0.005$ vs. P10.

### Supplementary Figure 6, Related to Fig. 5

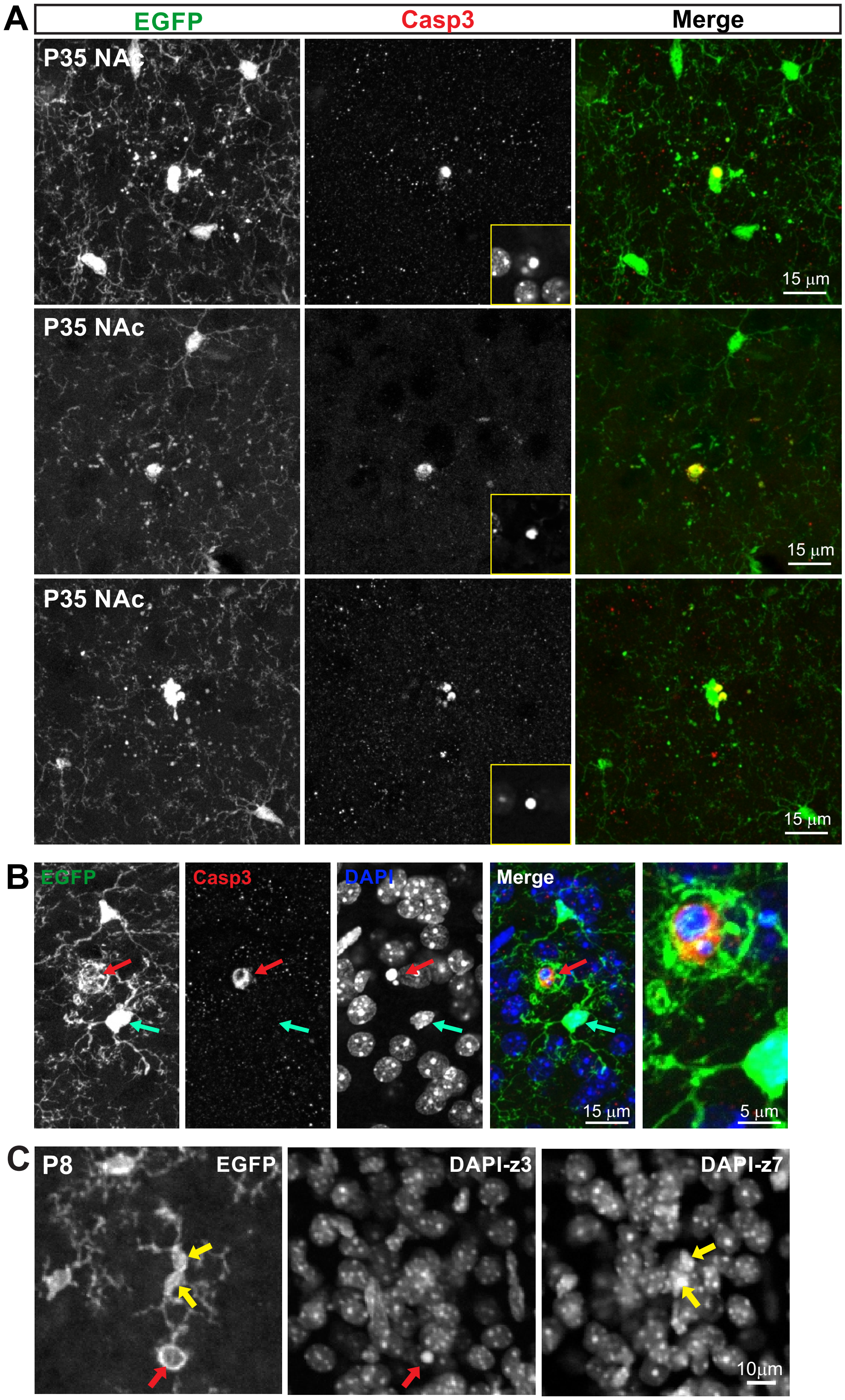

**Supplementary Figure 6. Microglial cells undergo programmed cell death during postnatal maturation. (A)** Additional examples of EGFP+Casp3+ microglia from NAc at P35. *Yellow box inset* shows DAPI labeling for the Casp3+ nucleus and indicates that all Casp3+ cells show highly condensed chromatin. (**B)** EGFP+Casp3+ microglia can clearly be distinguished from instances in which an EGFP+ microglial cell is engaging in phagocytotic engulfment of a neighboring Casp3+ cell. *Red arrow* highlights the Casp3+ nucleus and condensed chromatin of an EGFP- non-microglial cell. *Cyan arrow* highlights the absence of Casp3 labeling and the intact nucleus of the EGFP+ microglial cell engulfing the Casp3+ cell. Example shown is from P35 NAc. Not all z-levels for this max stack are shown in the DAPI channel for clarity. (**C**) Intact, dividing microglia phagocytose a nearby dying cell with condensed chromatin (*middle panel*). Nuclei of the dividing microglia (*right panel*) are not condensed.

### Supplementary Figure 7, Related to Fig. 7

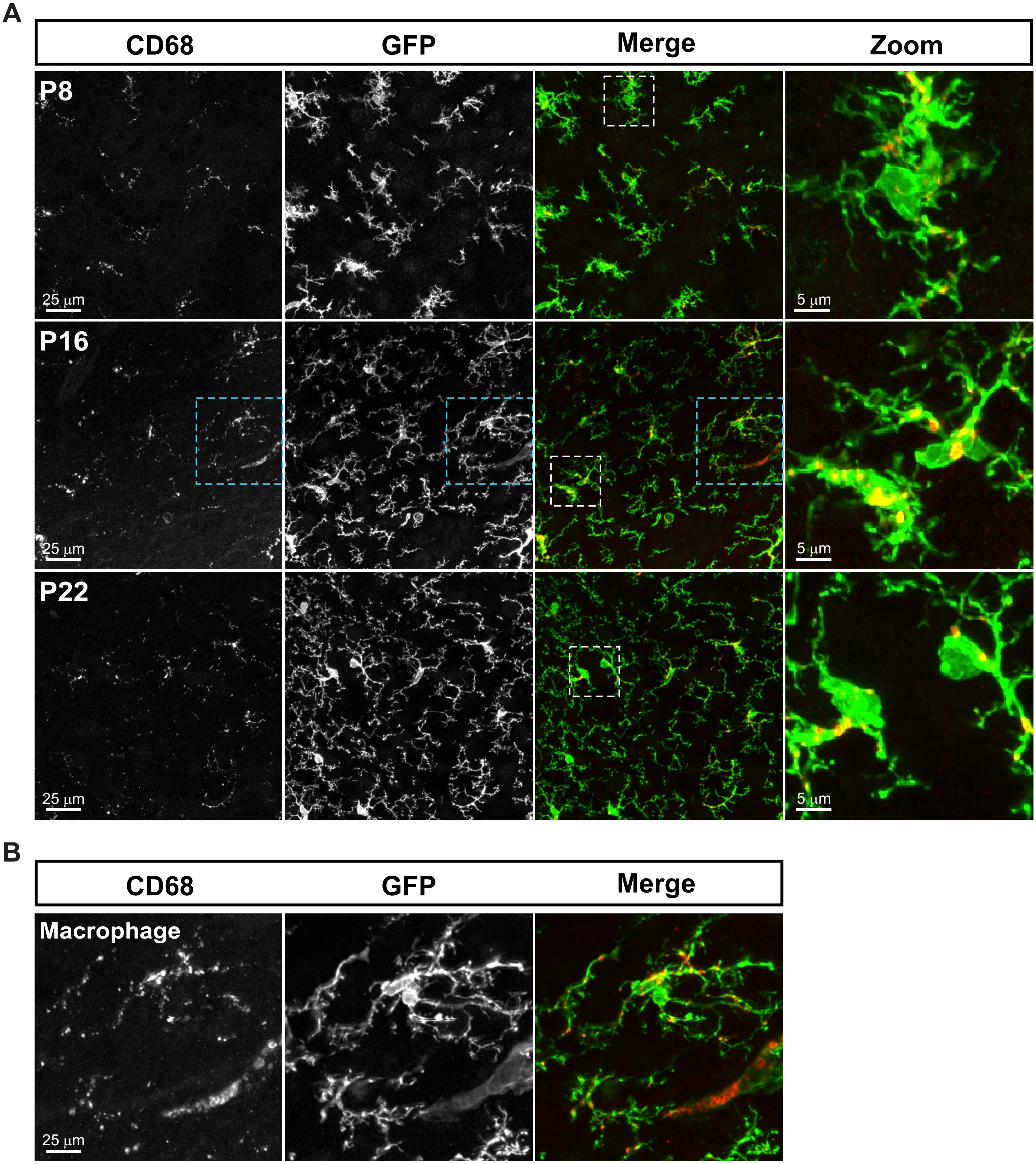

**Supplementary Figure 7. Microglial lysosome content within the VTA during early development.** **(A)** Representative images from the VTA of $CX3CR1^{EGFP/+}$ mice at different ages immunostained for Cluster of Differentation 68 (CD68) protein, a lysosome membrane protein expressed specifically in microglia and macrophages. Regions highlighted by *white boxes* shown at higher magnification *at right*. **(B)** Higher magnification view of the region highlighted by *blue boxes* in *A*. This field of view shows a microglial cell (*top middle*), and a macrophage (*bottom right*), highlighting the dimmer EGFP expression, lack of cell processes, and prominent CD68 labeling of macrophages. These cells could be unequivocally identified and removed from analysis.

### Supplementary Figure 8, Related to Fig. 2, 7, and S1

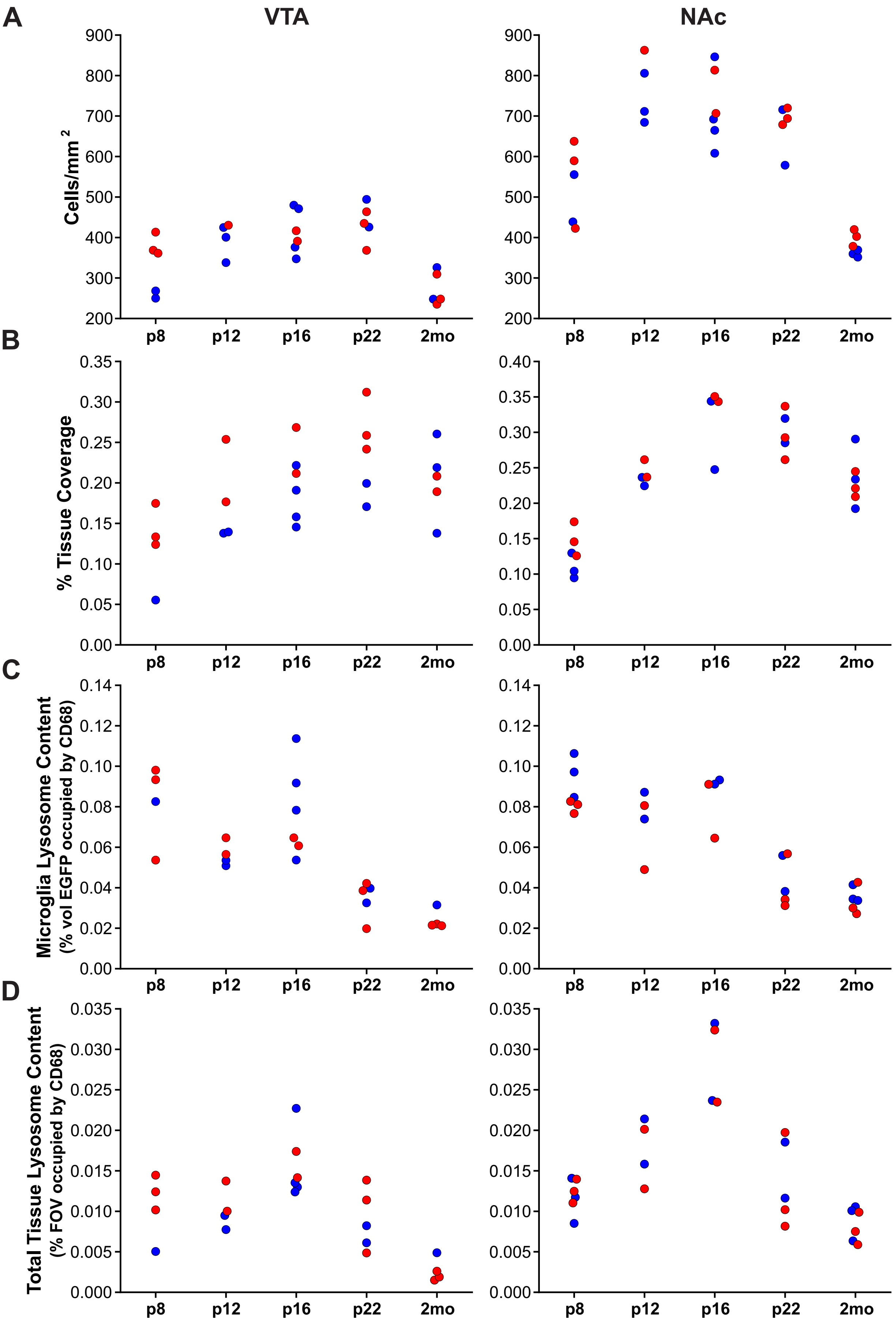

**Supplementary Figure 8. Absence of clear sexual dimorphism in analyzed parameters.** Graphs of main results colored according to the sex of the mouse (*male=blue, female=red*). **(A)** Quantification of microglial density from *CX3CR1*$^{EGFP/+}$ mice during early postnatal development in the VTA and NAc**. (B)** Quantification of microglia process coverage within the VTA and NAc during early postnatal development. **(C)** Quantification of microglial lysosome content during early postnatal development in the VTA and NAc. **(D)** Quantification of total tissue microglial lysosome content during early postnatal development in the VTA and NAc.
